# Exploring use of a protein cage system for producing bioactive peptides in *Escherichia coli*

**DOI:** 10.1101/2024.12.16.628579

**Authors:** Maxim D. Harding, Mark A. Jackson, Edward K. Gilding, Kuok Yap, David J. Craik, Frank Sainsbury, Nicole Lawrence

**Affiliations:** Institute for Molecular Bioscience, Australian Research Council Centre of Excellence for Innovations in Peptide and Protein Science, The University of Queensland, Brisbane, Queensland 4072, Australia; Centre for Cell Factories and Biopolymers, Institute for Biomedicine and Glycomics, Griffith University, Queensland 4111, Australia; Australian Research Council Centre of Excellence in Synthetic Biology, Australia

**Keywords:** bioactive peptide, recombinant expression, virus-like particle, protein encapsulation, bacteriophage P22

## Abstract

New therapeutics are urgently needed to curb the spread of drug-resistant diseases. Bioactive peptides (BAPs), including antimicrobial peptides, are emerging as an exciting new class of compounds with advantages over current drug modalities, especially small molecule drugs that are prone to resistance development. Here we evaluated a bacteriophage P22 virus-like particle (VLP) system where BAPs are encapsulated as fusion proteins with the P22 scaffold protein (SP) within self-assembling protein cages in *Escherichia coli*. Representative peptides from three structurally distinct classes of BAPs were successfully encapsulated into P22 VLPs at high cargo to VLP coat protein (CP) ratios that corresponded to interactions between the compact electropositive structures of the SP-BAPs and electronegative regions on the inward facing surface of CP subunits. However, high loading densities did not correspond to improved SP-BAP yields. An unexpected finding of this study was that while encapsulation alleviated negative effects of SP-BAPs on *E. coli* growth, the P22 scaffold protein acted as a sufficient fusion partner for accumulating BAPs, and co-expression of the CP did not further improve SP-BAP yields. Nevertheless, encapsulation in VLPs provided a useful first step in the purification pipeline for producing both linear and cyclic recombinant (r)BAPs that were functionally equivalent to their synthetic counterparts. Further efforts to optimise expression ratios of CP to SP-BAP fusions will be required to realise the full potential of encapsulation for protecting expression hosts and maximizing rBAP yields.

## Introduction

The threats caused by drug resistance are increasingly evident with the spread of antibiotic-resistant bacteria, multi-drug-resistant malaria parasites, and acquired drug-resistant cancers (Tacconelli, et al., 2018, Vasan, et al., 2019, Siqueira-Neto, et al., 2023). New therapeutics, and sustainable modes of producing them, are urgently needed. Antimicrobial peptides are a promising class of therapeutics that is gaining interest for treating these drug-resistant diseases. As this class comprises a diverse range of naturally occurring molecules with activities that extend into other diseases such as cancer, they can be more broadly defined as bioactive peptides (BAPs) (Magana, et al., 2020, Muttenthaler, et al., 2021). BAPs combine the advantages of small molecule drugs, such as membrane permeability and the potential for oral bioavailability, with those of larger biologic therapies such as antibodies, which have higher metabolic stability and target specificity (Craik, et al., 2013, Wang, et al., 2022). Moreover, resistance to peptides tends to occur more slowly, if at all, compared to small molecule drugs (Peschel and Sahl, 2006, Bell, 2011, Xuan, et al., 2023, Luo, et al., 2024). This reduced likelihood of inducing drug resistance may be due to peptides acting via multiple modes of action, but is most likely explained by the larger binding interfaces between BAPs and their molecular targets, which would require accumulation of multiple genetic mutations to disrupt interactions, in contrast to small molecule drugs which can be nullified by a single genetic mutation (Podlesek and Žgur Bertok, 2020, Amiss, et al., 2022). This benefit is exemplified by BAPs that kill target cells by membrane lysis, a mechanism of action that is not only difficult to overcome by acquired resistance, but which can also bypass established drug resistance mechanisms (Rodríguez-Rojas, et al., 2014, Benfield, et al., 2024).

Production of BAPs for lab-based discovery efforts and later for scaled industrial production has predominantly been achieved using synthetic chemistry approaches like solid-phase peptide synthesis (Wang, et al., 2022). Although chemical synthesis has enabled advances in peptide therapeutic development, excessive and harmful chemical waste from this approach has raised sustainability concerns that have triggered a movement for developing cheaper and greener production platforms (Wibowo and Zhao, 2019, Yap, et al., 2020, Roca-Pinilla, et al., 2022, Rossino, et al., 2023, Chaudhary, et al., 2024, Kekessie, et al., 2024). Challenges for producing recombinant (r)BAPs using greener recombinant expression platforms include their small size that makes them prone to proteolytic breakdown, and their biophysical properties such as high positive charge and amphipathic arrangement of charged and hydrophobic residues, which make them inherently toxic when they accumulate inside expression host cells (Li, 2011, Ishida, et al., 2016, Deo, et al., 2022). Applications of nanobiotechnology in recombinant production and metabolic engineering have recently expanded the biosynthetic range of production organisms for the manufacture of molecules of interest. For example, self-assembling protein cages have been used to compartmentalize and sequester recombinant products in *E. coli* to improve their stability and reduce toxicity (Patterson, et al., 2013, Das, et al., 2020, Lee, et al., 2020, Su, et al., 2023, Gawin, et al., 2025), an approach we here chose to explore for rBAP production.

Protein cages are nanoscale structures, comprised of one or more repeating protein subunits, that self-assemble to form a protein shell. Natural sources of protein cages include an array of independently evolved proteinaceous compartments, including viruses, ferritins, encapsulins, and bacterial microcompartments (Edwardson, et al., 2022). Recombinant protein cages also comprise engineered cages, the simplest being virus-like particles (VLPs) (McNeale, et al., 2023a), and a range of emerging *de novo* designed self-assembling protein compartments (Lutz, et al., 2023, Meador, et al., 2024, Yang, et al., 2024). Engineered protein cages have been used to encapsulate polypeptides such as enzymes for applications in metabolic engineering to produce valuable heterologous small molecules (Patterson, et al., 2014, Cheah, et al., 2021, Kang, et al., 2022, McNeale, et al., 2023b, Su, et al., 2023, Sator, et al., 2024), and shorter therapeutic peptides that require targeted delivery (Anand, et al., 2015, Wang, et al., 2018). In some cases, this approach has been shown to increase the yield of active proteins (Patterson, et al., 2013) or antimicrobial peptides (Lee, et al., 2020) in recombinant expression hosts. However, these examples have been demonstrated either only for a single peptide or have not successfully established downstream extraction protocols – probably due to a sub-optimal combination of encapsulating cage and respective recombinant cargo. Covalent attachment of cargo to protein cage coat proteins (CPs) to produce particles from single components can adversely affect assembly (Lee, et al., 2020, McNeale, et al., 2023a, Gawin, et al., 2025). Thus, two-component protein cage systems may be better suited for developing a modular BAP encapsulating system, as they allow better distinction between particle formation and cargo packaging.

Bacteriophage P22 VLPs form 58 nm icosahedral symmetries (T=7), comprising 420 CPs and ∼100-330 scaffold proteins (SPs) (Thuman-Commike, et al., 1998). Genetic fusion of heterologous proteins to either the N- or C-terminus of the SP, or a truncated version thereof, directs cargo packaging within the VLP through non-covalent interactions with the internal face of the CP (Parker, et al., 1998, O’Neil, et al., 2011). Recently, we developed a set of expression vectors to enable rapid testing of protein cargo encapsulation within P22 VLPs in *E. coli* (Esquirol, et al., 2022). Although P22 VLPs have primarily been applied for packaging enzymes for bio-catalysis, their ability to encapsulate heterologous proteins with fast packaging kinetics that ultimately shield these molecules from the host cell, makes them an ideal system for recombinant production of BAPs that are potentially toxic to expression host cells. Previous studies have demonstrated proof-of-concept towards this goal, for example with the rescue of otherwise insoluble proteins by encapsulation within P22 VLPs, and the recombinant packaging of BAPs such as the analgesic cone snail venom peptide MVIIA and the anticancer peptides NuBCP-9 and KLAK (Patterson, et al., 2013, Anand, et al., 2015, Wang, et al., 2018). However, in these studies the recombinant products were used in the context of the protein cage acting as a container for biocatalysis or a vehicle for therapeutic delivery, rather than as a manufacturing vessel from which the products could be successfully recovered.

Here, we used P22 VLPs in *E. coli* to produce positively charged rBAPs from three structurally distinct peptide classes: the horseshoe crab peptide tachyplesin I (TI; β-hairpin), which acts against bacteria (Edwards, et al., 2017, Amiss, et al., 2021) and its cyclic analog cTI, which acts against cancer cells (Vernen, et al., 2019, Benfield, et al., 2024); the mammalian defense peptide LL37 (α-helix), which exhibits both anticancer and antimicrobial activity (Dürr, et al., 2006, Kuroda, et al., 2015); and PDIP, a macrocyclic helix-loop-helix peptide engineered from the human defense protein platelet factor 4 with antimalarial (Lawrence, et al., 2018, Lawrence, et al., 2024) and anticancer (Lawrence, et al., 2020) activity. For PDIP, we also included a negatively charged linker between the positively charged SP binding domain and BAP sequences to demonstrate that decreasing overall negative charge and compactness of SP-BAP fusions adversely affects their packaging and assembly into P22 VLPs. Surprisingly, high packing densities of SP-BAP fusions did not translate to improved yield, which we attributed to incomplete encapsulation and the SP being able to act as a sufficient fusion partner for the BAPs in this study.

## Experimental Procedures

### Molecular cloning

Expression plasmids were cloned as previously described (Esquirol, et al., 2022). Briefly, gene blocks for all BAPs and mRUBY3 (control fluorescent reporter) were purchased from Integrated DNA Technologies in the form of dsDNA. Primers, designed with complementary BbsI sites were also ordered from Integrated DNA Technologies for Golden Gate cloning into the pRSFDuet expression vector, with CcdB-mediated counter selection. SP-BAP fusions were cloned in series with the P22 CP, and under control of the same T7 promoter. As a control to compare SP-fusion expression without encapsulation, pRSF constructs were also produced with the CP removed. Each of the constructs was transformed into Top 10 *E. coli* cells, with clones selected in the presence of 50 µg/mL kanamycin. The correct identity of cloned fragments was confirmed by sequencing (Australian Genome Research Facility).

### Expression and purification of P22 VLP encapsulated SP-BAP fusions

Transformed cells were grown in LB broth supplemented with 50 µg/mL kanamycin. Expression cultures were induced with 1 mM final concentration IPTG and incubated for 16 h at 28 °C. Following overnight expression, cells were pelleted by centrifugation at 4,500 g for 15 min at 4°C. Cell pellets were resuspended in phosphate buffered saline (PBS, pH 7.4) before three passages through a cell press at 25 kpsi. Cell lysates were clarified by centrifugation at 25,000 g for 20 min at 4 °C. Clarified protein extract was then loaded onto an iodixanol density gradient (15, 20, 25%) and centrifuged at 150,000 g for 3 h at 4 °C using a Beckman Coulter L-100XP floor standing ultracentrifuge. To isolate encapsulated SP-BAP fusions, VLPs were extracted from the bottom of ultracentrifuge tubes and desalted into PBS using a PD-10 column. VLP concentrations were calculated using a Pierce BCA assay (ThermoFisher Scientific), then analyzed by SDS-PAGE and tryptic digest LC-MS/MS as above. To determine cargo loading, protein bands from SDS-PAGE were quantified using ImageJ (Esquirol, et al., 2022) and compared to determine the ratio of CP and SP fusion cargo. These densitometric calculations were also used in combination with the BCA assay data to calculate the recombinant yield of SP-BAP fusions as mg/L of culture media.

### Modeling of SP-BAP structure and interaction with CP

The structure of P22 CP, assembly into interacting subunits, and interactions between CP and the binding domain of SP were determined from PDB: 8I1V (Xiao, et al., 2023). The structures of SP-fused to PDIP (including with EEG linkers), TI, or LL37 were modeled with AlphaFold2 on ColabFold v1.5.5. Interactions between the binding domain of SP fused BAPs (including EEG linkers for PDIP) and CP were modeled with AlphaFold2-multimer with MMseqs2 on ColabFold v 1.5.5 (see Table S2 for truncated input sequences for SP-BAPs).

The model used to predict the structure of monomers (SP-BAP) was alphafold2_ptm. For SP-BAP:CP interactions the alphafold2_multimer_v3 model was used. All runs comprised three recycles, a maximum of 200 iterations, used a greedy pairing strategy, and had five output 3D structures. The highest ranked structure was selected to represent the most likely structure of each SP-BAP or truncated SP-BAP:CP interaction.

All structure diagrams were produced using PyMOL Molecular Graphics System, v 2.5.2 Schrodinger, LLC. Electrostatic potential surface diagrams were prepared by using PDB2PQR and APBS software (https://server.poissonboltzmann.org) (Jurrus, et al., 2018) to generate dx input files and then performing molecular surface visualization using the APBS Electrostatics plugin in PyMOL.

### Negative stain Transmission Emission Microscopy (TEM)

Purified VLPs were diluted to 0.25 mg/mL in PBS and applied (10 µL) to formvar/carbon coated grids (ProSciTech) for 2 min. Grids were washed three times on droplets of water for 30 seconds before staining on 10 µL of 2% uranyl acetate for 2 min. TEM was performed on a JEOL 1010 electron microscope at 80 kV operating voltage.

### Growth of *E. coli* expressing encapsulated compared to free SP-BAP fusions

Growth of *E. coli* expressing encapsulated SP-BAP (co-expression with CP) was compared to free SP-BAP expressed from CP-deficient constructs. Starter cultures were grown in LB broth with 50 µg/mL kanamycin at 37 °C, with shaking at 220 rpm, until the bacteria were in growth phase (OD_600_ between 0.4 and 0.8). Cultures were diluted to an OD_600_ of 0.1 in LB broth with 50 µg/mL kanamycin, induced with IPTG (1 mM final concentration), and plated into 96 well plates (100 µL per well). The plate was incubated at 28 °C in a TECAN SPARK plate reader, and A_600_ was measured every 30 min for 16 h. Growth curves were prepared from the averages of two technical repeats for each of three biological replicates. Crude extracts of *E. coli* total cell protein collected at 16 h were run on SDS-PAGE to compare SP-BAP yields as a percentage of the total protein content.

### Extraction and processing of rBAPs

VLPs purified by ultracentrifugation and desalted into PBS (approximately 3.5 mL) were diluted into 50 mL of chaotropic cargo extraction buffer (6.75 M urea, 500 mM NaCl, 50 mM Tris, 20 mM imidazole, pH 7.4) and shaken over night at 4 °C. The disassembled VLP solution was loaded onto a HisTrap FF column (Cytiva) and SP-BAPs were purified from the P22 CP using an ÄKTA FPLC system. Samples were analyzed by SDS-PAGE following purification before buffer exchange using a PD10 column into tobacco etch virus (TEV) protease processing buffer (50 mM Tris, 1 mM DTT, 0.5 mM EDTA, pH 8.0). Buffer-exchanged SP-BAPs were then concentrated using a 3.5 kDa Amicon filter (Merck) before being quantified using a Pierce BCA assay (ThermoFisher Scientific). rBAPs were then cleaved from the SP by overnight incubation with TEV enzyme variant hyperTEV60 (Sumida, et al., 2024) at 4 °C at a 1:10 (enzyme:substrate) molar ratio. Success of cleavage was assessed both by SDS-PAGE and on a Sciex 5800 matrix-assisted laser desorption/ionization (MALDI) TOF-MS. Cleaved rBAPs were then purified from the reaction by reverse-phase high-performance liquid chromatography (RP-HPLC) using a Shimadzu system and a Phenomenex Jupiter C18 column with a 1% gradient from solvent A (0.05% (v/v) TFA in water) to solvent B (0.05% (v/v) TFA in 90% acetonitrile). HPLC fractions were assessed by electro-spray ionization mass spectrometry (ESI-MS) and the rBAP-containing fractions were lyophilized. For cyclisation to produce rcTI and rcPDIP, the lyophilized linear peptides were resuspended in cyclisation buffer (100 mM Na_3_PO_4_ and 50 mM NaCl at pH 6) and incubated with modified recombinant *Oldenlandia affinis* asparaginyl endopeptidase (AEP) enzyme ([C247A]*Oa*AEP1b), as previously described (Yap, et al., 2021). Briefly, AEP enzyme and rBAP substrate were incubated overnight at room temperature at a ratio of 1:10 (enzyme:substrate). Success of the overnight reaction was confirmed by MALDI TOF-MS as described above.

### Antimicrobial activity of purified rRAPS

*E. coli* (ATCC 25922) was grown in LB broth to mid-log growth phase (as above) and diluted to 8 x 10^5^ colony forming units per mL (A_600_ = 0.001). Peptide treatments were prepared in LB and serially diluted into 96-well plates, which were incubated for 18 h at 37 °C following addition of diluted bacteria. Minimum inhibitory concentration (MIC) values were determined from wells where no bacterial growth was observed (verified by measuring A_600_) for two biological replicates. Synthetic peptides were prepared as before (Benfield, et al., 2024). Wells with no treatment or 50 µM gentamicin were included as controls for 0% and 100% inhibition, respectively.

## Results and Discussion

### Protein cage construct design

With the aim of developing a reliable platform for the biosynthetic production of a range of BAPs in *E. coli*, we investigated the effect of encapsulation within P22 VLPs on rBAP yield. We used the recently developed P22 plasmid system (Esquirol, et al., 2022) that enables Golden Gate cloning of desired genes into the two-component pRSFDuet plasmid such that the coding sequences are introduced in-frame with the P22 SP, fused at either the N- or C-terminus (Supplementary Table S1), to be co-expressed with the P22 CP (Figure 1).

**Figure 1.**
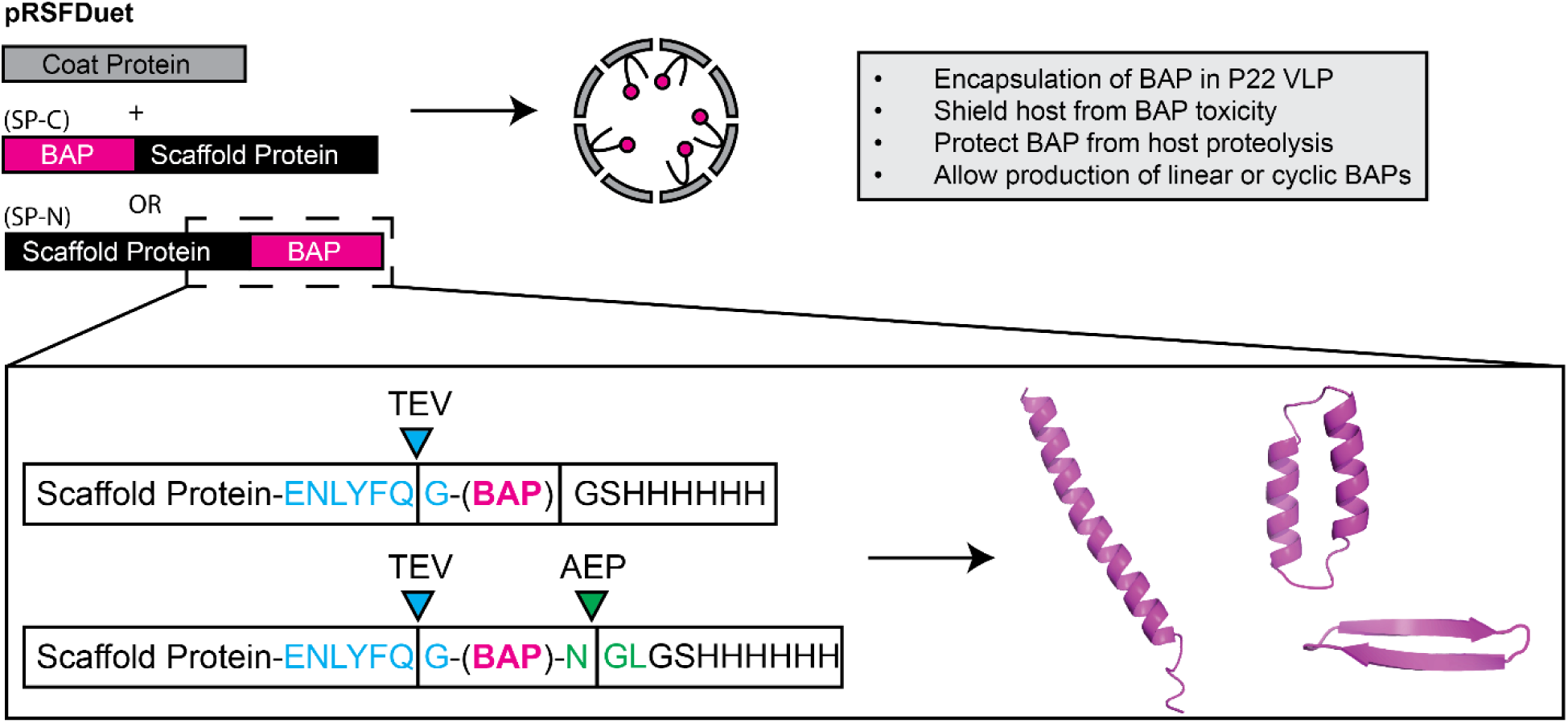
Protein cage construct design. pRSFDuet expression constructs were designed to co-express the SP-BAP fusion (either SP-N or SP-C configuration) and the P22 CP, with the aim of packaging BAPs into VLPs to shield the *E. coli* host from toxic BAP effects and the BAP from host proteases. SP-BAP fusions were designed such that a range of structurally distinct rBAPs could be extracted and processed from the VLP by protease cleavage *in vitro*. The tobacco etch virus (TEV) protease recognition sequence (blue) allows liberation of recombinant (r)BAPs from the SP fusion partner. The *Oldenlandia affinis* asparaginyl endopeptidase (AEP; [C247A]*Oa*AEP1b) recognition sequence (green) provides a site for simultaneous cleavage of the (His)6 purification tag and peptide bond formation between N- and C-terminal rBAP residues.

The pRSFDuet constructs were designed to enable straightforward purification and processing steps for downstream liberation of encapsulated SP-BAP fusions from P22 VLPs. A hexa-histidine tag was included at either the N- or C-termini, whichever was distal to the SP among the different constructs to enable downstream chromatographic separation of the SP-BAP fusion from the P22 CP. To facilitate cleavage of rBAPs from the SP-fusion, a tobacco etch virus (TEV) protease processing site was included immediately adjacent to the BAP N-terminus, enabling the cleaved rBAP to retain only a single glycine (G) amino acid scar at its N-terminus. To enable peptide backbone cyclisation – a modification employed to enhance stability and resistance to proteolytic breakdown of therapeutic peptides (Chan, et al., 2013, Craik and Du, 2017, Vernen, et al., 2019, Hayes, et al., 2021) – an asparaginyl endopeptidase (AEP) recognition sequence was included at the BAP C-terminus to enable subsequent cyclisation (Harris, et al., 2015, Yang, et al., 2017).

### Evaluating the expression and packaging of BAPs into P22 VLPs

To determine whether the P22 VLP is a suitable system for biosynthetic production of BAPs, we assembled constructs with the therapeutic antimalarial peptide candidate PDIP and the P22 SP fused to either the N-terminus or C-terminus (SP-N, SP-C; Figure 1). As a control, we produced constructs with mRUBY3 as a fluorescent reporter of expression. Using the pRSFDuet vectors to co-express SP-mRUBY3 or SP-PDIP together with P22 CP allowed successful expression and encapsulation of both mRUBY3 and PDIP, as observed by SDS-PAGE of purified VLPs for both SP-N and SP-C fusions (Figure 2C and 2D, respectively). Correct identity of the SP-fusions and P22 CP was confirmed using tryptic digest LC-MS/MS (Supplementary Table S2). This is the first time that the antimalarial peptide candidate PDIP has been successfully expressed in any format by *E. coli*. Packaging of SP-PDIP fusions produced heterogeneous VLPs with some particles having altered morphology (Figure 2E). Correctly formed VLPs have 420 CP, and it is customary to report cargo loading in terms of the number of SP-fusion molecules per particle. To account for possible variation from 420 CP per particle, here we instead report loading density as SP-fusion (cargo) to CP ratio. Both SP-N and SP-C orientations of SP-PDIP fusion proteins had higher cargo to CP ratios (1.4, 1.7) than SP-mRUBY3, which had cargo to CP ratios of 0.6 for both SP-N and SP-C fusions, see Figure 2B.

**Figure 2.**
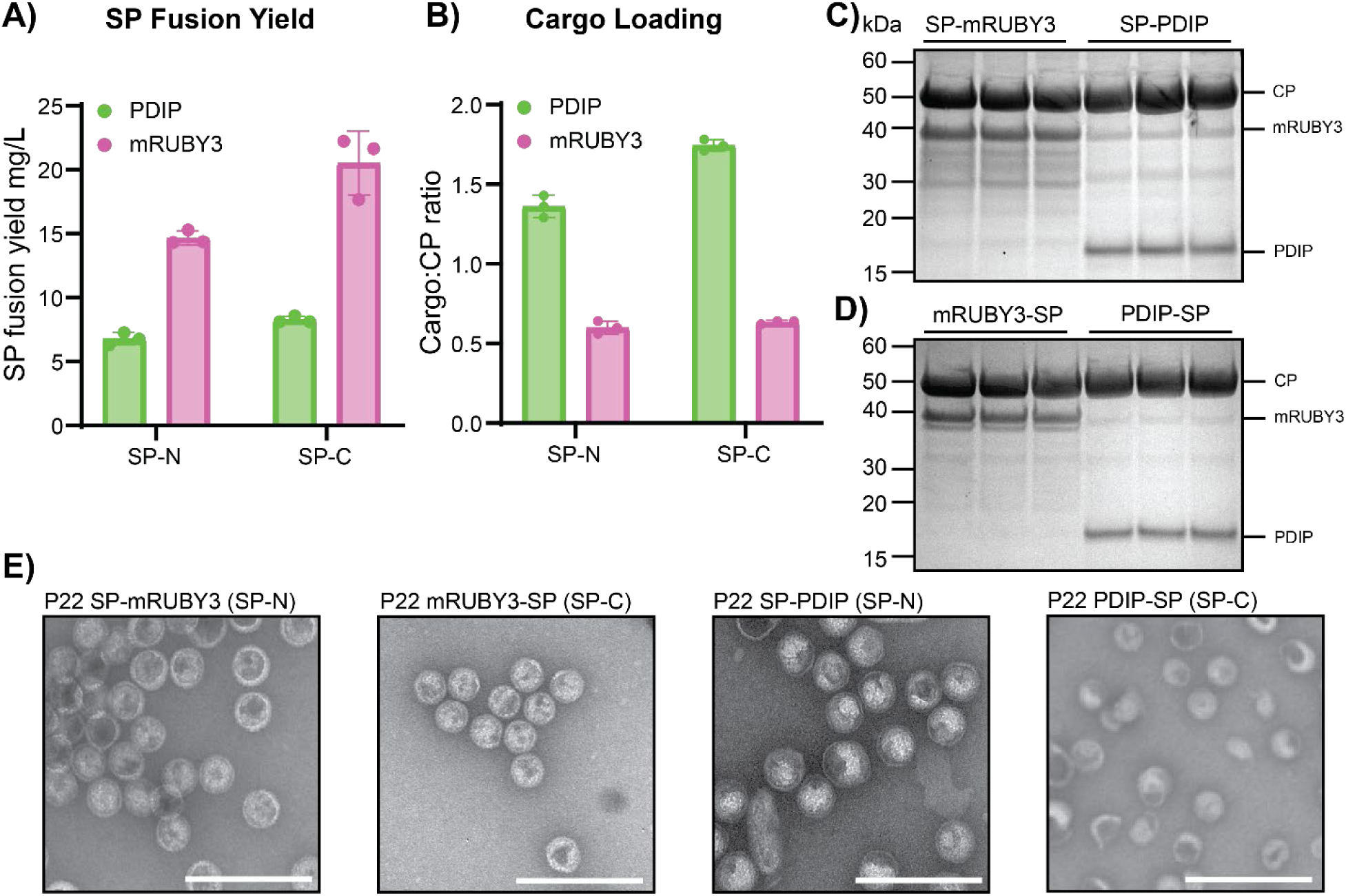
Evaluation of P22 VLPs for packaging PDIP and mRUBY in SP-N and SP-C orientations. A) Comparison of recombinant yields of VLP-associated SP-fusions (mg/L) expressed in SP-N or SP-C orientations. B) Calculated SP-fusion cargo to CP ratio. C) SDS-PAGE for SP-N expressions for mRUBY3 and PDIP. D) SDS-PAGE for SP-C expressions for mRUBY3 and PDIP. Data in panels A-D are from three separate expressions for each construct. E) Representative negative stain TEM images for all four prototype P22 VLPs. The white scale bar is 250 nm.

The high observed packaging ratio of PDIP as both SP-N and SP-C fusions suggests favorable interactions between P22 CP and SP-fusion. This finding is consistent with the high cargo to CP packaging density of 1.3 (543 cargo fusion proteins per VLP, assuming 420 CP per correctly formed particle) that was previously reported for the analgesic peptide MVIIA, which was packaged to explore delivery across the blood brain barrier (Anand, et al., 2015).

Encapsulation of the fusion proteins allowed production of PDIP-SP fusions at yields of 6.7 mg/L for SP-N, 8.2 mg/L for SP-C, and mRUBY3 SP fusions at yields of 14.3 mg/L for SP-N, 20.5 mg/L for SP-C (Figure 2A). Assessment of the purified particles by negative stain transmission electron microscopy (TEM) confirmed the successful assembly of P22 VLPs for each of the four prototype SP fusions (Figure 2E). Compared to purified particles packaged with mRUBY3, the VLPs with packaged PDIP displayed irregularity in particle morphology for both SP-N and SP-C orientations (Figure 2E, Supplementary Figure S1). This irregularity might be caused by the high ratio of SP-fusion to CP, which can lead to excess assembly nuclei, aberrant assembly, and formation of kinetically trapped assembly intermediates (Parent, et al., 2006). Despite the observed particle heterogeneity, the capacity to recombinantly express PDIP in this system suggests that the *E. coli* host cells have been protected to some extent from PDIP’s membrane-disruptive activity.

Given the minor difference in recombinant yield achieved for encapsulated PDIP in either the SP-N or SP-C orientation (Figure 2A–D), further work here used the SP-N orientation only, for two reasons. First, for linear BAPs such as LL37, removal of the SP via TEV cleavage from the SP-N orientation leaves a single glycine residue on the final product (Figure 1). Second, we found that AEP processing (cyclization) of PDIP was inefficient when using the SP-C orientation (Supplementary Figure S2), possibly due to the intrinsically disordered nature of the N-terminal region of the SP in solution (Guo, et al., 2024), which may interfere with enzyme binding or activity (McNeale, et al., 2023b).

### SP-PDIP binding interactions and VLP packaging

To gain a better understanding of potential binding interactions between SP-PDIP and the P22 CP, we compared the previously reported interactions between the structured C-terminal region of the SP and an internal facing region of the CP (Xiao, et al., 2023) (Figure 3A) and the predicted binding interaction (AlphaFold2 MMseqs2) between SP-PDIP and the CP. SP-PDIP is predicted to form a compact structure with the helix-loop-helix structures of each component contributing to an electropositive patch (Figure 3B). Overlay of the predicted interaction between CP and SP-PDIP, and the reported structures of CP and SP binding domains in Figure 3C suggests that the SP component of the fusion binds CP in a near identical location to free SP, whereas the PDIP component binds additional adjacent sites (CP) with negative electrostatic potential on the inner face of the VLP.

**Figure 3.**
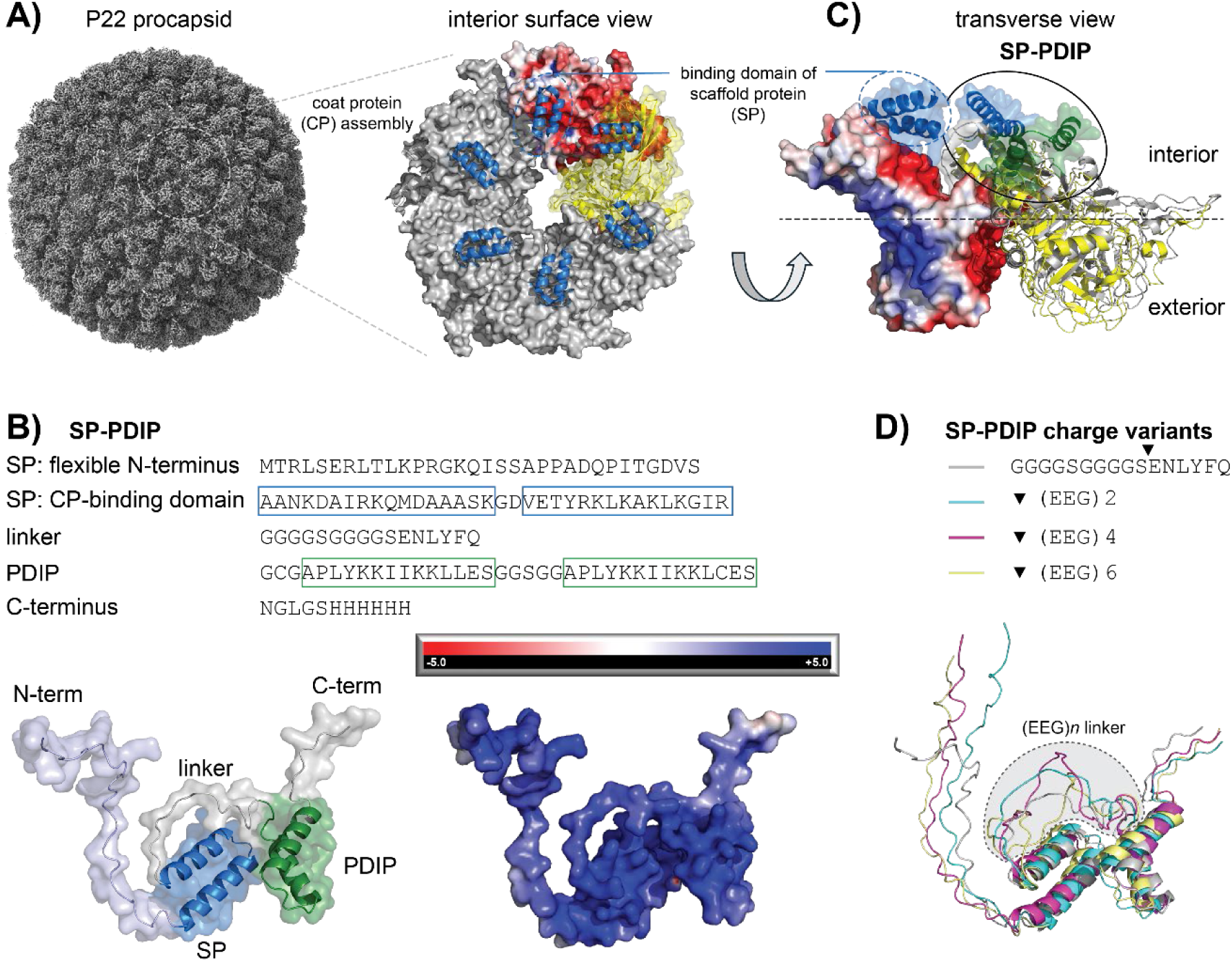
Predicted interaction of SP-PDIP on the internal face of P22 VLPs. **A**) The P22 procapsid (X-axis view of 3D surface, EMD-35124) is formed from repeating assemblies of coat proteins (CPs). The inner surface has electronegative regions that bind to, and are stabilized by, electropositive regions at the C-terminus of scaffold proteins (SPs), shown as blue helix-loop-helix cartoons. The electrostatic surface potential is shown for one of the CP subunits with the red color indicating electronegative potential. Structures and interactions were determined from PDB:8I1V. **B**) Amino acid sequence and predicted structure of SP-PDIP (AlphaFold2) with helix-loop-helix structures of the CP-binding domain of SP (blue) and PDIP (green) associating into a compact electronegative patch, as indicated by the blue color on the electrostatic surface potential diagram (right). **C**) Transverse view showing two CP subunits (left, electrostatic surface potential; right yellow ribbon) and bound SP (blue transparent surface with ribbon) (from PDB:8I1V), overlaid with the predicted structures and interaction of CP (grey ribbon) with SP-PDIP (blue+green transparent surface with ribbon) (AlphaFold2-multimer with MMseqs2). **D**) Predicted structures (AlphaFold2) of SP-PDIP and charge variants with inserted [EEG]*n* linkers between SP C-terminus and PDIP N-terminus. All structure diagrams were produced using PyMOL v 2.5.2. Visualization of electrostatic surface potential was performed using input files from PDB2PQR and APBS software (Jurrus, et al., 2018).

### Influence of charge on BAP packaging efficiency

The considerable difference in cargo loading between PDIP and mRUBY3 (Figure 2B) led us to investigate contributing factors. We hypothesized that in addition to its small size compared to mRUBY3 and other previously reported proteins (O’Neil, et al., 2011), the high positive charge of PDIP promotes higher cargo loading. This would make the P22 VLP encapsulation system, with its negatively charged interior surface, particularly well-suited for packaging molecules with positively-charged surfaces, a common feature of many BAPs. To test this, we designed a series of charge-varied SP-PDIP constructs that incorporated anionic linker sequences, comprised of 2, 4, or 6 repeating motifs of the amino acids glutamic acid and glycine (EEG) between the C-terminal CP-binding domain of SP and PDIP. We used the approach of including anionic linkers because it has been successfully employed to mask the activity of adjacent positively charged antimicrobial peptides, including TI (Ngambenjawong, et al., 2022).

The three SP-PDIP variants, together with the original SP-PDIP construct, providing a series of with an overall charge ranging from +13 to +1 (Figure 4A, 4B), were expressed from the pRSFDuet plasmid to assess the influence of overall charge on P22 VLP loading efficiency. Each of the charge-varied constructs was successfully packaged into P22 VLPs as shown by negative stain TEM (Figure 4C), although the heterogenous mixtures with numerous aberrant particles suggested that the (EEG)*n* linkers may affect VLP assembly. The expressed SP-(EEG)*n*-PDIP constructs were visualised by SDS-PAGE of purified VLPs (Figure 4D, Supplementary Figure S3) and correct full-length sequences were confirmed by tryptic digest LC-MS/MS (Supplementary Table S2). Densitometric analysis of the cargo loading for each charge variant, revealed a sequential decrease in cargo to CP ratio (1.4, 1.2, 1.0, 0.5) for SP-PDIP followed by the charge variants with 2, 4, then 6 EEG, that corresponded to a decrease in overall charge (Figure 4E).

**Figure 4.**
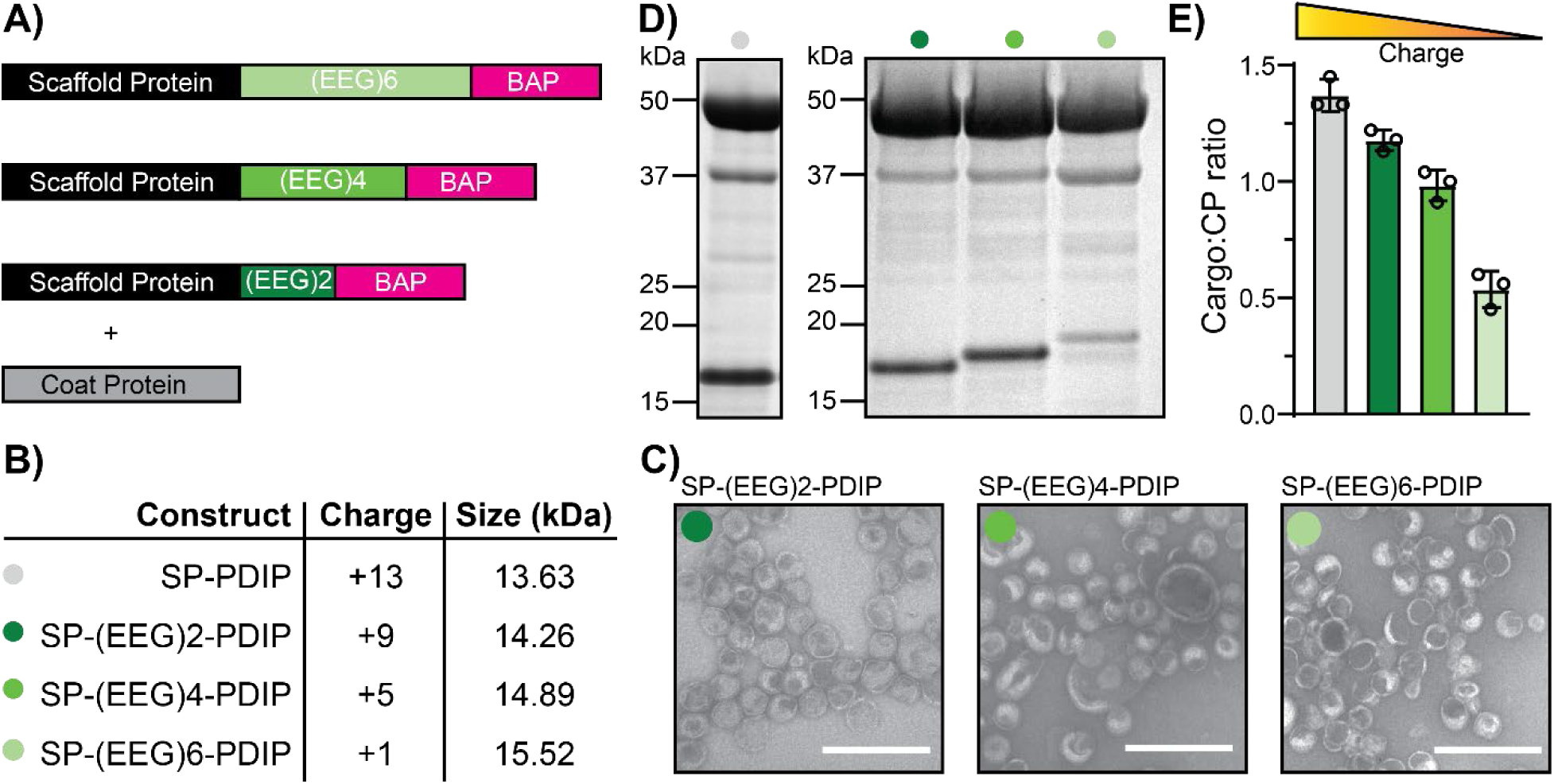
Effect of charge on cargo packaging. A) Constructs for anionic linker series of SP-PDIP fusions. B) The calculated charge of the constructs at pH 7.4 and size in kDa. Colour code for each construct is shown alongside construct names. C) Negative stain TEM images for SP-(EEG)2-PDIP, SP-(EEG)4-PDIP, and SP-(EEG)6-PDIP from left to right respectively. White scale bar represents 250 nm. D) SDS-PAGE image for comparison of SP-PDIP band size and intensity with anionic linker changes. E) Calculated cargo to CP ratio from n = 3 biological replicates.

It is possible that the sequential increase in length of the (EEG)*n* linker contributed to the decrease in packaging density due to increased steric hinderance of SP:CP, PDIP:CP, or CP:CP interactions during VLP assembly. However, the increase in size is small (650 Da for each (EEG)2 added) compared to the size of each SP-(EEG)*n*-PDIP fusion (ranging from 4-12% total protein fusion mass for (EEG)2-(EEG)6. Furthermore, structure predictions suggest that the helix-loop-helix structure of both the C-terminal binding domain of SP and PDIP are maintained irrespective of the length of the central (EEG)*n* linker (Figure 4D). Thus, we propose it is more likely that reduction of the overall positive charge of the cargo is responsible for this trend for several reasons. First, using the same expression vectors, we recently reported SP-fusion cargo to CP loadings as high as 1.4 (588 cargo per 420 CP, or one correctly formed VLP) for a 55 kDa enzyme (McNeale, et al., 2023b). In that case, high loading was attributed to oligomerisation of the enzymes, but it does show that loading density is not necessarily restricted by increased cargo bulk. Second, previous work investigating P22 SP-CP interactions and cargo retention in P22 VLPs showed that a truncated version of the P22 SP (the SP sequence used in this study) had greater affinity for the P22 CP compared to the larger wild-type SP (McCoy, et al., 2018). This increased affinity was also attributed to increased overall positive charge, rather than decreased size, of the truncated SP compared to the wild-type. These combined observations suggest that the small compact structures and high positive charge of SP-BAP fusions promote increased loading within P22 VLPs and that this platform is well suited for expressing and accumulating a range of cationic BAPs.

### Expression of two structurally different cationic BAPs using P22 VLPs

To assess the broader applicability of the P22 protein cage system for biosynthetic production of BAPs, we tested the expression of BAPs from two additional structural classes; the linear α-helical peptide LL37 (Sancho-Vaello, et al., 2020) and a cyclic analogue of the β-hairpin peptide TI (cTI; Vernen, et al., 2019, Benfield, et al., 2024). Like SP-PDIP, SP-TI and SP-LL37 are predicted to have a compact structure, and to bind to electronegative regions of CP subunits that face the inside of VLPs (Supplementary Figure S4). Co-expression of TI and LL37 as SP-N fusions with P22 CP (pRSFDuet system) successfully facilitated P22 VLP assembly and BAP cargo encapsulation, as shown by SDS-PAGE, negative stain TEM, and tryptic digest LC/MS-MS (Figure 4A, 4B, and Table S2). Despite the apparent heterogeneity of the resultant VLPs with some particles showing an aberrant morphology (Figure 5A, B), yields for encapsulated SP-LL37 and SP-TI were 12.3 and 3.3 mg/L of culture respectively (Figure 5C), with high density cargo to CP loading ratios of 1.5 for SP-LL37 and 2.4 for SP-TI (Figure 5D).

**Figure 5.**
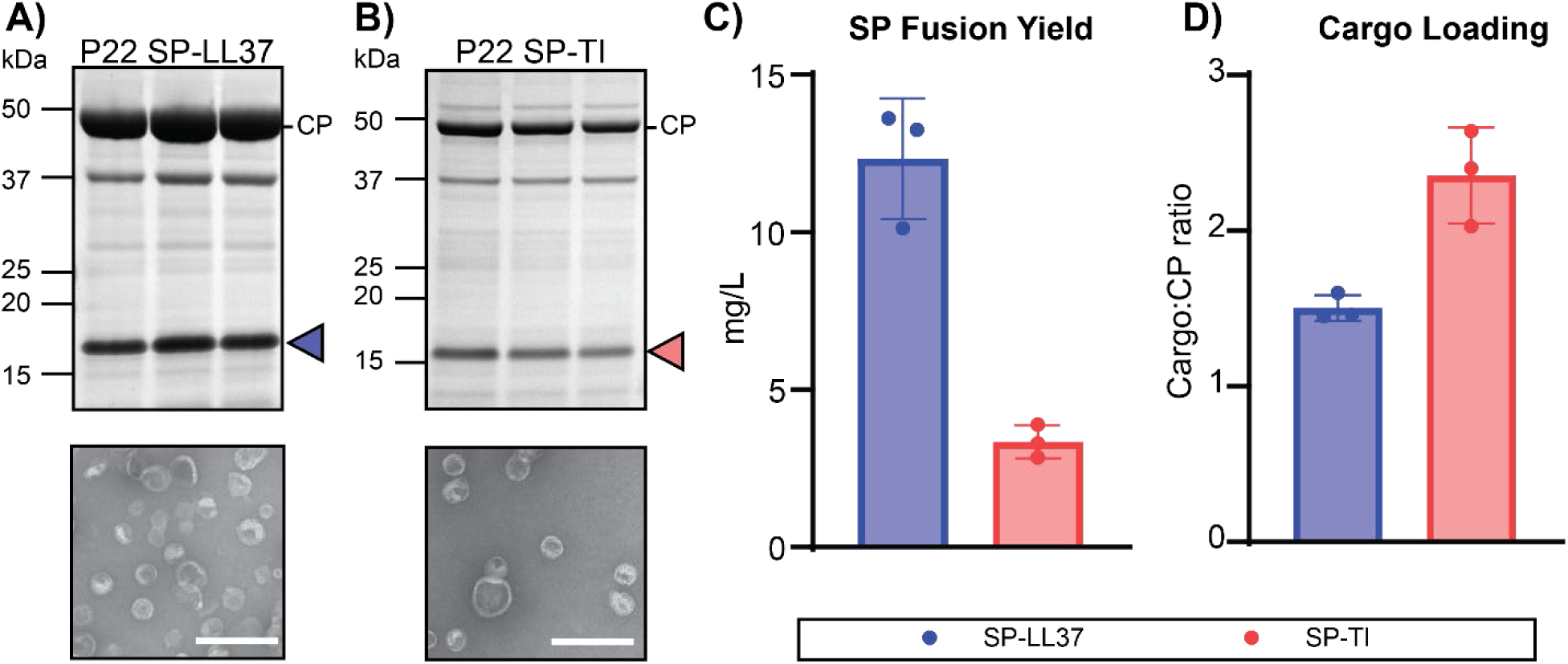
Encapsulation of LL37 and TI as SP fusions in P22 VLPs. A) SDS-PAGE and negative stain TEM of P22 SP-LL37. Blue arrow indicates SP-LL37. The scale bar on TEM image indicates 250 nm. B) SDS-PAGE and negative stain TEM of P22 SP-TI. Red arrow indicates SP-TI. The scale bar on TEM image indicates 250 nm. C) SP-BAP fusion yields as mg/L of culture for SP-LL37 and SP-TI expressed using pRSFDuet SP-N (n = 3 biological replicates). D) Calculated SP-fusion cargo to CP ratio for P22 SP-LL37 (blue) and SP-TI (red).

The cargo to CP loading density of SP-TI was higher than observed for SP-PDIP and SP-LL37 despite these three SP-BAPs having a similar overall charge. It is possible that the more compact secondary structure (β-sheet) of TI (see Supplementary Figure S4) allows better interaction with electronegative regions on CP subunits to facilitate greater packaging for SP-TI than the helical SP-BAP fusions. However, this high cargo loading for SP-TI was not reflected in the yield (Figure 5C), as SP-TI had the lowest yield of all BAPs tested. TI is a potent broad spectrum antimicrobial peptide, with killing mechanisms that involve membrane permeabilizaton and also binding to DNA and RNA (Hong, et al., 2015, Edwards, et al., 2017). One possible explanation for the inverse relationship between the high apparent cargo loading but comparatively low yield of SP-TI is that if not completely packaged, the more potent antimicrobial activity of TI (Edwards, et al., 2017, Amiss, et al., 2021) may impact cell viability and reduce overal recombinant protein production.

### Effect of encapsulation on *E. coli* growth and SP-BAP production

To further interrogate our hypothesis that encapsulation of BAPs within P22 VLPs will shield host *E. coli* cells from toxic effects of the BAPs and/or shield the BAPs from proteolytic breakdown, we compared the growth of *E. coli* transformed with the pRSFDuet constructs that coexpress SP-BAP and CP (encapsulated SP-BAP) to those transformed with control pRSF constructs with the CP removed (free SP-BAP). Analysis of growth curves over 16 h revealed a disparity in growth kinetics (Supplementary Figure S5) between induced (+IPTG) and uninduced (-IPTG) *E. coli*. Expression of free SP-fusions reduced *E. coli* growth for all tested constructs, including SP-mRUBY3. In comparison, expression of encapsulated SP-constructs had a less pronounced and differential effect on growth compared to uninduced *E. coli.* Here, co-expression of the CP (encapsulation) almost completely overcame growth inhibition by SP-mRUBY, and partially improved *E. coli* growth for SP-PDIP, SP-(EEG)_4_-PDIP, and SP-LL37. SP-TI was the exception, with similar growth inhibition observed for free compared to encapsulated fusion proteins. (Supplementary Figure S5). These observations suggest that co-expression of the P22 CP and consequent VLP encapsulation does improve *E. coli* growth when expressing SP-BAPs. However, assessment of total protein content after 16 h induction did not demonstrate an increase in SP-BAP yield for encapsulated compared to unencapsulated expressions (Supplementary Figure S6). These unexpected outcomes suggest that P22 SP was a sufficient fusion partner for expressing and accumulating the BAPs tested in this study and that encapsulation into VLPs did not provide any additional advantage in terms of SP-fusion yield.

The following additional observations were made from comparing the SP-fusion to CP ratio from the total cell proteins (TCP) and VLP-associated proteins (see Figure S7). For all constructs, there was a higher proportion of SP-fusion in the TCP. This observation suggests that not all cargo was packaged into VLPs. Therefore, production of two disparately sized proteins (46.7 kDa CP compared to 12–15 kDa SP-BAP) from a single construct did not produce ideal ratios for complete cargo encapsulation. Interestingly, the amount of available SP-fusion (Figure S6, S7) did not explain differences in cargo loading density (VLP). This is especially evident when comparing SP-PDIP and the SP-(EEG)4-PDIP charge variant, which supports a role of differential charge-driven interactions with CP subunits rather than differences in expression levels, in determining cargo packing density.

### Processing and purification of recombinant (r)BAPs

Despite our observation that encapsulation within P22 VLPs did not improve the yield of SP-fusions, it still provides a valuable first step in the purification pipeline, whereby VLP-associated proteins (CP and SP-fusions) can be isolated from *E. coli* proteins using ultracentrifugation. Thus we employed a processing pipeline with four steps: (1) purify P22 VLPs containing SP-BAPs from total soluble protein by ultracentrifugation; (2) isolate SP-BAPs from VLPs using IMAC under chaotropic conditions that dissassemble the P22 protein cage (Capen and Teschke, 2000); (3) enzymatically cleave rBAPs from SP using TEV protease; (4) purify liberated rBAPs using HPLC (Figure 6A). An additional step can be included for BAPs that require further processing, for example backbone cyclization to produce cyclic peptides.

**Figure 6.**
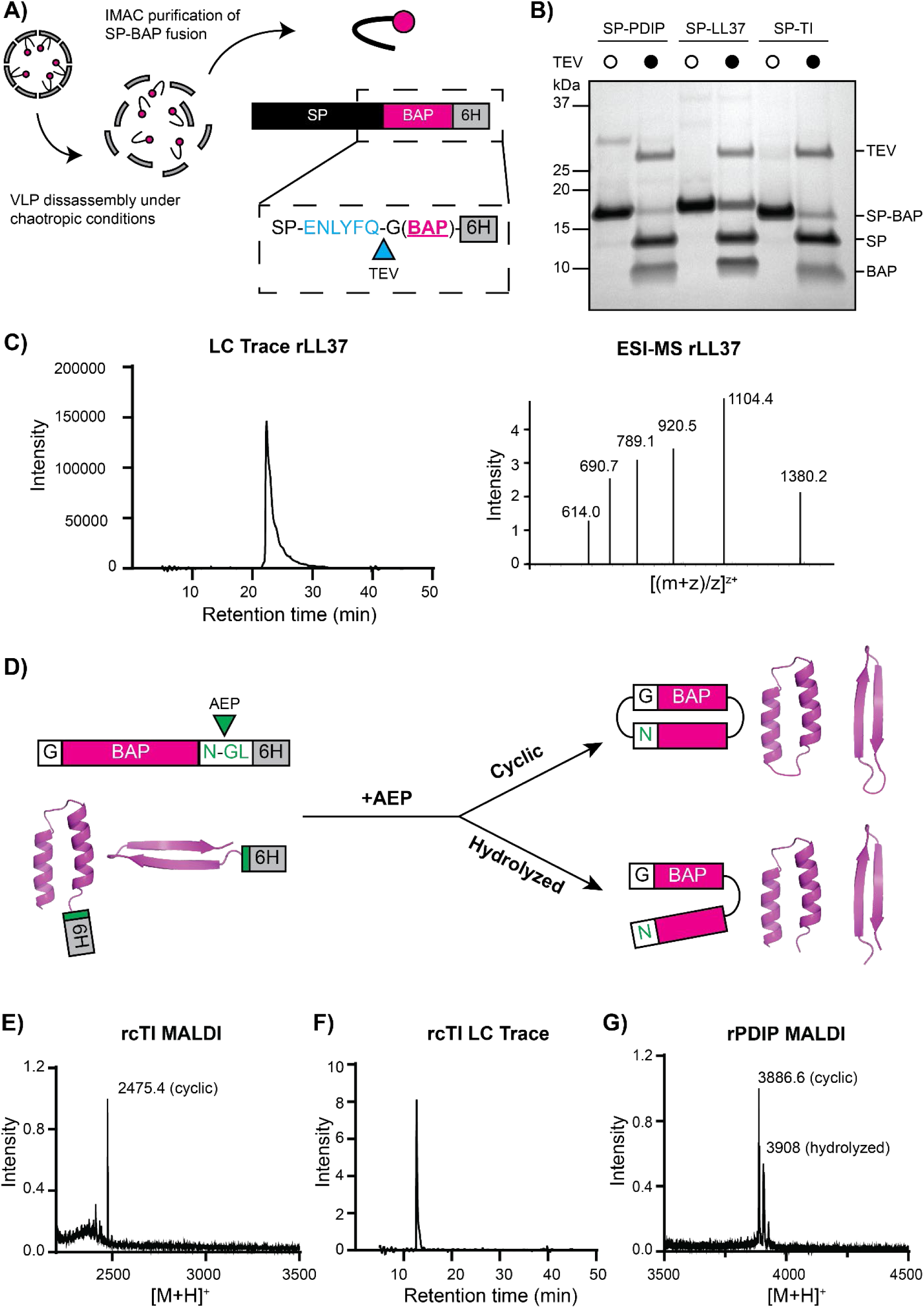
Post-expression processing of rBAPs from P22 VLPs. A) Illustration showing VLP disassembly to release SP-BAPs. Each SP-BAP has a TEV recognition site (blue), an N-terminal glycine (required for cyclization reactions), and a C-terminal hexa-histidine tag. B) SDS-PAGE image of TEV (hTEV60)-mediated SP-BAP cleavage for all three BAPs tested. Empty white circles above lanes indicate no added enzyme and black circles indicate addition of TEV. Bands are labelled on the right side of gel. C) LC trace of cleaved rLL37 purified from the TEV cleavage reaction by HPLC and the corresponding MS signal from LC-MS. D) Illustration of AEP processing reaction. Addition of AEP to purified rBAPs containing a C-terminal NGL recognition sequence can lead to one of two products: the correctly processed desired cyclic product, or a hydrolyzed linear version. E) MALDI-MS spectra of the AEP ([C247A]*Oa*AEP1b) reaction for production of rcTI. F) LC trace for rcTI. G) MALDI-MS spectra of the AEP ([C247A]*Oa*AEP1b) reaction to produce rcPDIP (with both cyclic and hydrolyzed forms detected).

To confirm the technical feasibility of this processing pipeline, we tested the purification and processing of all three BAPs that were expressed as SP-fusions and encapsulated within P22 VLPs in this study. Under chaotropic conditions, IMAC efficiently separated all tested SP-BAPs from P22 VLPs via binding of the C-terminal hexa-histidine tag (Supplementary Figure S8). Following buffer exchange of purified SP-BAPs into TEV processing buffer, overnight incubation with hTEV60 (Sumida, et al., 2024) successfully liberated each of the rBAPs from their SP fusion partner (Figure 6B). Subjecting these TEV-cleavage reactions to HPLC successfully enabled purification of the cleaved rBAPs, as characterized by LC-MS (Supplementary Figures S9, S10, and S11). This is the final processing step required for rLL37, and for other linear BAPs that could be expressed using this system, as the LC trace and correct MS peaks confirmed the isolation of pure rLL37 with a C-terminal hexa-histidine tag (Figure 6C). Notably, re-arrangement of the expression vector to place the histidine tag at the N-terminus of the SP would enable production of linear rBAPs, with a minimal N-terminal glycine residue remaining after TEV cleavage.

Cyclization of rPDIP and rTI was achieved by overnight incubation of the purified linear peptides (containing C-terminal AEP recognition motif NGL) with [C247A]*Oa*AEP1b enzyme at a 1:10 enzyme:substrate ratio (Yang, et al., 2017, Yap, et al., 2021) (Figure 6D). Analysis of the reaction using MADLI-TOF MS revealed complete processing of rTI to produce recombinant cyclic TI (rcTI) (Figure 6E), which was successfully purified using HPLC and characterized with LC-MS (Figure 6F, Supplementary Figure S12). However, MALDI-MS analysis of the rPDIP cyclization reaction revealed the presence of both cyclic (rcPDIP) and hydrolyzed forms (Figure 6G), that were unable to be separated using HPLC. This outcome is consistent with our attempts to cyclize other PDIP analogues that were generated via chemical synthesis. Therefore, enzyme-mediated cyclization of PDIP needs further optimization of conditions to improve the yield of rcPDIP compared to hydrolyzed or unprocessed linear peptide when using AEP enzymes. Alternatively, other enzymes might provide a superior outcome for cyclizing PDIP. For example, sortase A has been used to successfully cyclize disulfide rich peptides (Jia, et al., 2014) and has even been included as a fusion partner in biosynthetic pathways to generate self-cyclizing peptides (Jia, et al., 2023).

### Antimicrobial activity of rBAPs

An important requirement for recombinantly produced BAPs is that they have similar activity to their synthetically derived counterparts. Thus, we determined the minimum inhibitory concentration (MIC) of rcTI and rLL37 compared to cTI and LL37 that were produced using solid phase peptide synthesis methodologies, as previously described (Benfield, et al., 2024). Serially diluted peptides were prepared in LB media then added to growth phase *E. coli* (ATCC 25922) at 8 x 10^5^ colony forming units/mL (A_600_ = 0.001) in 96-well plates. After 16 h incubation at 37 °C, MIC were identified from wells with no visible growth, which was verified by measuring A_600_ and comparing to wells with 0% (untreated) and 100% (50 µM gentamicin) growth inhibition. Despite the small changes in amino acid sequence between recombinant and synthetic BAPs – additional Gly and Asn for rcTI, and N-terminal Gly, C-terminal Gly-Ser-(His)6 for rLL37 – the rBAPs were at least as active against *E. coli* as their synthetic counterparts (Table S4). The MICs observed in the current study are higher than previously reported for cTI (1 µM (Vernen, et al., 2019)) and LL37 (30 µg/mL, equivalent to ∼7 µM (Aghazadeh, et al., 2019)), but this difference is most likely due to the current assays being performed in LB (to provide direct comparison to the growth media used for expression of rBAPs) instead of Mueller Hinton Broth (a more optimized media for determining MIC using microbroth dilution methods)(CLSI, 2024).

## Conclusions

In this study we used the two-component bacteriophage P22 system, with SP-fusions (cargo) and CP expressing from a single high copy pRSFDuet vector in *E. coli*, to produce positively charged BAPs rLL37, rcTI, and rPDIP. This is the first time that PDIP, and to our knowledge cTI, have been produced biosynthetically. Heterogeneous populations of VLPs were produced for each of the BAPs, with high observed SP-fusion to CP packaging ratios that were most likely due to interactions between the compact electropositive surfaces of the SP-BAPs and inward-facing electronegative regions of CP subunits. Indeed, the packaging ratio for SP-PDIP was decreased by including anionic linkers between SP and PDIP, suggesting that this approach might provide a useful strategy for tuning cargo loading.

Surprisingly, the high SP-fusion to CP packaging ratios did not translate to an increased yield compared to expression of the SP-BAP fusions alone. This could possibly be due to an insufficient supply of CP relative to the SP-BAP, causing malformed VLP assemblies and a failure to achieve optimal cargo accumulation. The key effect of packaging density on yield of polypeptides produced using protein cage encapsulation has recently been demonstrated using the *Aquifex aeolicus* lumazine synthase (AaLS) cage system (Gawin, et al., 2025). That study employed a two-component packaging approach (separate vectors with CP and CP-fusion) to increase recombinant yields in *E. coli* by reducing the relative packaging of therapeutic peptide candidates, including the AMP magainine-2, within AaLS cages. Similarly, the P22 VLP system could be modified to optimize particle formation compared to SP-cargo packaging, by expressing the CP and SP-BAPs from different plasmids. For example, expressing the SP-BAPs with a weaker promoter and/or from a lower copy number plasmid may prevent overaccumulation of the smaller cargo (12–15 kDa) compared to the CP (46.7 kDa). Additionally, optimizing growth conditions could be explored to overcome the plateau in bacterial growth that was observed when the CP was co-expressed with the SP-BAP fusions.

This study highlights the importance of validating all aspects of cargo encapsulation, including the effects of expressing multiple proteins on *E. coli* growth, the influence of cargo properties (including charge) on encapsulation and particle formation, and the fate of co-expressed cargo. For the BAPs included in this study, we observed that the SP acted as a sufficient expression partner to allow accumulation in *E. coli* hosts, whereas co-expression with CP to encapsulate SP-BAPs improved *E. coli* growth, but not the observed yield compared to total cell protein. The mechanism of protection afforded by the SP fusion partner remains unclear, but it is most likely due to the formation of insoluble aggregates, similar to previous reports for TI (Panteleev, et al., 2017). By contrast, here we demonstrated that encapsulation in P22 VLPs provided a means of isolating soluble VLP-associated proteins from *E. coli* total cell protein, which served as an excellent starting point for a pipeline to produce linear and cyclic BAPs from three structurally different classes. For the cyclic peptides, incubation of linear rTI with AEP ligase resulted in complete conversion to rcTI, which was readily purified. However, pure rcPDIP could not be isolated from a hydrolyzed reaction by-product using this approach, suggesting the need for alternative cyclization strategies for that peptide. Accordingly, the P22 VLP encapsulation system could provide an alternative pathway to solid-phase peptide synthesis, that reduces input costs and environmental burden (Kekessie, et al., 2024). Given the P22 plasmid system can accommodate co-loading of different cargoes *in vivo* (McNeale, et al., 2023b), co-encapsulation of SP-BAP fusions and downstream processing enzymes (e.g. TEV, AEP, sortase A) could be explored to minimize *in vitro* processing requirements. With many therapeutic BAPs employing non-proteinogenic amino acids, future work could apply this P22 VLP platform in genetically expanded *E. coli* lines capable of incorporating such amino acids into BAP sequences.

## Author Contributions

**Maxim D. Harding** conceptualization, data curation, formal analysis, investigation, methodology, project administration, validation, visualization, writing – original draft; **Mark A. Jackson** funding acquisition, methodology, resources, supervision, writing – review and editing; **Edward K. Gilding** funding acquisition, methodology, supervision, writing – review and editing; **Kuok Yap** methodology, writing – review and editing; **David J. Craik** funding acquisition, resources, supervision, writing – review and editing; **Frank Sainsbury** conceptualization, methodology, resources, validation, writing – review and editing; **Nicole Lawrence** funding acquisition, methodology, project administration, resources, supervision, validation, writing – review and editing.

## Acknowledgements

This work was supported by funding from the US Department of Defense Defense Congressionally Directed Medical Research Program (PR210354 to DJC, NL, MAJ, EKG) and the Australian Research Council Centre of Excellence for Innovations in Peptide and Protein Science (CE200100012). DJC was supported by NHMRC grant (2009564). FS is supported by an Australian Research Council Future Fellowship (FT230100084) and the Australian Research Council Centre of Excellence in Synthetic Biology (CE200100029).

