## Supplementary Information for "Exploring use of a protein cage system for producing bioactive peptides in *Escherichia coli*"

**Contents**

Table S1. Amino acid sequences of proteins used in this study

Table S2. Tryptic digest LC-MS/MS of VLP-associated proteins expressed in this study

Table S3. Truncated input sequences for modelling interaction between SP-BAPs and CP

Table S4. Comparison of *E. coli* MIC for recombinant cTI and LL37 and their synthetic counterparts

Figure S1. Expanded TEM images of representative P22 VLPs expressed in this study

Figure S2. Inefficient AEP processing of SP-C PDIP fusion

Figure S3. SDS-PAGE gel of VLP-associated proteins from SP-(EEG)*n*-PDIP expression

Figure S4. Predicted interactions between SP-BAP fusions and P22 CP

Figure S5. Growth curves of *E. coli* expressing encapsulated (pRSFDuet) compared to free (pRSF) SP-fusions

Figure S6. Comparison of total proteins from *E. coli* expressing encapsulated (pRSFDuet) compared to free (pRSF) SP-fusions

Figure S7. Comparison of cargo to coat protein (CP) ratio for SP-fusions

Figure S8. Immobilized metal affinity chromatography (IMAC) and SDS-PAGE of extracted SP-BAP fusion proteins

Figure S9. HPLC traces for purification of rBAPs after TEV cleavage

Figure S10. UPLC trace of TEV cleaved rBAPs

Figure S11. ESI-MS of TEV-cleaved and HPLC-purified rBAPs

Figure S12. LC-MS of rcTI

#### Table S1. Amino acid sequences of proteins used in this study [a]

| Protein name & molecular weight | Amino acid sequence |
| --- | --- |
| P22 CP<br>46.7 kDa | MALNEGQIVTLAVDEIIETISAITPMAQKAKKYTPPAASMQRSSNTIWMPVEQESPTQEGW<br>DLTDKATGLLELNVAVNMGEPDNDFFQLRADDLRDETAYRRRIQSAARKLANNVELKVA<br>NMAAEMGSLVITSPDAIGTNTADAWNFVADAEEIMFSRELNRDMGTSYFFNPQDYKKAG<br>YDLTKRDIFGRIPEEAYRDGTIQRQVAGFDDVLRSPKLPVLTKSTATGITVSGAQSFKPVAW<br>QLDNDGNKVNVDNRFATVTLSATTGMKRGDKISFAGVKFLGQMAKNVLAQDATFSVVR<br>VVDGTHVEITPKPVALDDVSLSPEQRAYANVNTSLADAMAVNILNVKDARTNVFWADDA<br>IRIVSQPIPANHELFAGMKTTSFSIPDVGLNGIFATQGDISTLSGLCRIALWYGVNATRPEAIG<br>VGLPGQTA |
| SP-PDIP<br>13.6 kDa | MTRLSERLTLKPRGKQISSAPPADQPITGDVSAANKDAIRKQMDAAASKGDVETYRKLKA<br>KLKGIRGGGGSGGGGS*ENLYFQG*CGAPLYKKIIKKLLESGGSGGAPLYKKIIKKLCES**NGL**<br>GSHHHHHH |
| PDIP-SP<br>13.6 kDa | MHHHHHHGS*ENLYFQG*CGAPLYKKIIKKLLESGGSGGAPLYKKIIKKLCES**NGL**GSGGGG<br>SGGGGTRLSERLTLKPRGKQISSAPPADQPITGDVSAANKDAIRKQMDAAASKGDVETYR<br>KLKAKLKGIR |
| SP-mRUBY3<br>36.1 kDa | MTRLSERLTLKPRGKQISSAPPADQPITGDVSAANKDAIRKQMDAAASKGDVETYRKLKA<br>KLKGIRGGGGSGGGGSMVSKGEELIKENMRMKVVMEGSVNGHQFKCTGEGEGRPYEG<br>VQTMRIKVIEGGPLPFAFDILATSFMYGSRTFIKYPADIPDFFKQSFPEGFTWERVTRYEDG<br>GVVTVTQDTSLEDGELVYNVKVRGVNFPSNGPVMQKKTKGWEPNTEMMYPADGGLRG<br>YTDIALKVDGGGHLHCNFVTTYRSKKTVGNIKMPGVHAVDHRLERIEESDNETYVVQRE<br>VAVAKYSNLGGGMDELYKGSHHHHHH |
| mRUBY3-SP<br>35.3 kDa | MHHHHHHGSMVSKGEELIKENMRMKVVMEGSVNGHQFKCTGEGEGRPYEGVQTMRIK<br>VIEGGPLPFAFDILATSFMYGSRTFIKYPADIPDFFKQSFPEGFTWERVTRYEDGGVVTVTQ<br>DTSLEDGELVYNVKVRGVNFPSNGPVMQKKTKGWEPNTEMMYPADGGLRGYTDIALKV<br>DGGGHLHCNFVTTYRSKKTVGNIKMPGVHAVDHRLERIEESDNETYVVQREVAVAKYSN<br>LGGGMDELYKGSGGGGSGGGGTRLSERLTLKPRGKQISSAPPADQPITGDVSAANKDAIR<br>KQMDAAASKGDVETYRKLKAKLKGIR |
| SP-LL37<br>14.1 kDa | MTRLSERLTLKPRGKQISSAPPADQPITGDVSAANKDAIRKQMDAAASKGDVETYRKLKA<br>KLKGIRGGGGSGGGGS*ENLYFQG*LLGDFFRKSKEKIGKEFKRIVQRIKDFLRNLVPRTESG<br>SHHHHHH |
| SP-TI<br>12.2 kDa | MTRLSERLTLKPRGKQISSAPPADQPITGDVSAANKDAIRKQMDAAASKGDVETYRKLKA<br>KLKGIRGGGGSGGGGS*ENLYFQG*KWCFRVCYRGICYRRCRG**NGL**GSHHHHHH |
| SP-EEG2-PDIP<br>14.2 kDa | MTRLSERLTLKPRGKQISSAPPADQPITGDVSAANKDAIRKQMDAAASKGDVETYRKLKA<br>KLKGIRGGGGSGGGGSEEGEEG*ENLYFQG*CGAPLYKKIIKKLLESGGSGGAPLYKKIIKKL<br>CES**NGL**GSHHHHHH |
| SP-EEG4-PDIP<br>14.9 kDa | MTRLSERLTLKPRGKQISSAPPADQPITGDVSAANKDAIRKQMDAAASKGDVETYRKLKA<br>KLKGIRGGGGSGGGGSEEGEEGEEGEEG*ENLYFQG*CGAPLYKKIIKKLLESGGSGGAPLYK<br>KIIKKLCES**NGL**GSHHHHHH |
| SP-EEG6-PDIP<br>15.5 kDa | MTRLSERLTLKPRGKQISSAPPADQPITGDVSAANKDAIRKQMDAAASKGDVETYRKLKA<br>KLKGIRGGGGSGGGGSEEGEEGEEGEEGEEGEEG*ENLYFQG*CGAPLYKKIIKKLLESGGSG<br>GAPLYKKIIKKLCES**NGL**GSHHHHHH |

[a] CP, P22 coat protein; SP, P22 scaffold protein; PDIP, Platelet factor 4 derived internalization peptide; mRUBY3, fluorescent reporter protein; LL37, human cathlecidin; TI, tachyplesin I. Specific protein motifs are coded as follows. Hexa-histidine tag for nickel affinity chromatography of cargo proteins is shown in red. The BAP sequences are shown in blue. Processing site for tobacco etch virus protease (TEV) are *underlined and italicised*. Processing site for asparaginyl endopeptidase (AEP) is **underlined and bolded**. Anionic block linkers are shown in green for the charge altered SP-PDIP constructs.

#### Table S2. Tryptic digest LC-MS/MS of VLP-associated proteins expressed in this study [a]

| Protein name | Amino acid sequence |
|---|---|
| P22 CP | MALNEGQIVTLAVDEIIETISAITPMAQKAKKYTPPAASMQRSSNTIWMPVEQESPTQEGWDLTDKATGLLELNVAVNMGEPDNDFFQLRADDLRDETAYRRRIQSAARKLANNVELKVANMAAEMGSLVITSPDAIGTNTADAWNFVADAEEIMFSRELNRDMGTSYFFNPQDYKKAGYDLTKRDIFGRIPEEAYRDGTIQRQVAGFDDVLRSPKLPVLTKSTATGITVSGAQSFKPVAWQLDNDGNKVNVDNRFATVTLSATTGMKRGDKISFAGVKFLGQMAKNVLAQDATFSVVRVVDGTHVEITPKPVALDDVSLSPEQRAYANVNTSLADAMAVNILNVKDARTNVFWADDAIRIVSQPIPANHELFAGMKTTSFSIPDVGLNGIFATQGDISTLSGLCRIALWYGVNATRPEAIGVGLPGQTA |
| SP-PDIP | MTRLSERLTLKPRGKQISSAPPADQPITGDVSAANKDAIRKQMDAAASKGDVETYRKLKAKLKGIRGGGGSGGGGSENLYFQGCGAPLYKKIIKKLLESGGSGGAPLYKKIIKKLCESNGLGSHHHHHH |
| SP-mRUBY3 | MTRLSERLTLKPRGKQISSAPPADQPITGDVSAANKDAIRKQMDAAASKGDVETYRKLKAKLKGIRGGGGSGGGGSMVSKGEELIKENMRMKVVMEGSVNGHQFKCTGEGEGRPYEGVQTMRIKVIEGGPLPFAFDILATSFMYGSRTFIKYPADIPDFFKQSFPEGFTWERVTRYEDGGVVTVTQDTSLEDGELVYNVKVRGVNFPSNGPVMQKKTKGWEPNTEMMYPADGGLRGYTDIALKVDGGGHLHCNFVTTYRSKKTVGNIKMPGVHAVDHRLERIEESDNETYVVQREVAVAKYSNLGGGMDELYKGSHHHHHH |
| SP-LL37 | MTRLSERLTLKPRGKQISSAPPADQPITGDVSAANKDAIRKQMDAAASKGDVETYRKLKAKLKGIRGGGGSGGGGSENLYFQGLLGDFFRKSKEKIGKEFKRIVQRIKDFLRNLVPRTESGSHHHHHH |
| SP-TI | MTRLSERLTLKPRGKQISSAPPADQPITGDVSAANKDAIRKQMDAAASKGDVETYRKLKAKLKGIRGGGGSGGGGSENLYFQGKWCFRVCYRGICYRRCRGNGLGSHHHHHH |
| SP-EEG2-PDIP | MTRLSERLTLKPRGKQISSAPPADQPITGDVSAANKDAIRKQMDAAASKGDVETYRKLKAKLKGIRGGGGSGGGGSEEGEEGENLYFQGCGAPLYKKIIKKLLESGGSGGAPLYKKIIKKLCESNGLGSHHHHHH |
| SP-EEG4-PDIP | MTRLSERLTLKPRGKQISSAPPADQPITGDVSAANKDAIRKQMDAAASKGDVETYRKLKAKLKGIRGGGGSGGGGSEEGEEGEEGEEGENLYFQGCGAPLYKKIIKKLLESGGSGGAPLYKKIIKKLCESNGLGSHHHHHH |
| SP-EEG6-PDIP | MTRLSERLTLKPRGKQISSAPPADQPITGDVSAANKDAIRKQMDAAASKGDVETYRKLKAKLKGIRGGGGSGGGGSEEGEEGEEGEEGEEGEEGENLYFQGCGAPLYKKIIKKLLESGGSGGAPLYKKIIKKLCESNGLGSHHHHHH |

[a] Protein species from the analysed SDS-PAGE shown in Figure 2, Figure 4 and Figure 5 were identified by an in-gel trypsin digest of excised bands followed by LC-MS/MS analysis. Underlined letters indicate peptide fragments identified by in gel tryptic digest LC-MS/MS.

#### Table S3. Truncated input sequences for modelling interaction between SP-BAPs and CP[a]

| Component/s | Amino acid sequence |
|---|---|
| SP-PDIP | AANKDAIRKQMDAAASKGDVETYRKLKAKLKGIRGGGGSGGGGS*ENLYFQG*CGAPLYKKIIKKLLESGGSGGAPLYKKIIKKLCES |
| SP-EEG2-PDIP | AANKDAIRKQMDAAASKGDVETYRKLKAKLKGIRGGGGSGGGGSEEGEEG*ENLYFQG*CGAPLYKKIIKKLLESGGSGGAPLYKKIIKKLCES |
| SP-EEG4-PDIP | AANKDAIRKQMDAAASKGDVETYRKLKAKLKGIRGGGGSGGGGSEEGEEGEEGEEG*ENLYFQG*CGAPLYKKIIKKLLESGGSGGAPLYKKIIKKLCES |
| SP-EEG6-PDIP | AANKDAIRKQMDAAASKGDVETYRKLKAKLKGIRGGGGSGGGGSEEGEEGEEGEEGEEGEEG*ENLYFQG*CGAPLYKKIIKKLLESGGSGGAPLYKKIIKKLCES |
| SP-TI | AANKDAIRKQMDAAASKGDVETYRKLKAKLKGIRGGGGSGGGGS*ENLYFQG*KWCFRVCYRGICYRRCRG |
| SP-LL37 | AANKDAIRKQMDAAASKGDVETYRKLKAKLKGIRGGGGSGGGGS*ENLYFQG*LLGDFFRKSKEKIGKEFKRIVQRIKDFLRNLVPRTES |

[a] SP-BAP sequences were truncated to remove the unstructured N-terminal region of SP (determined from PDB:8I1V) and the C-terminus of the fusion protein comprising the AEP processing site and 6xHis tag. The truncated SP is shown in black, adjoining TEV cleavage site in underlined italics, BAP sequence in blue, and additional (EEG)n linker in green.

#### Table S4. Comparison of *E. coli* MIC for recombinant cTI and LL37 and their synthetic counterparts

| | Origin | Amino acid sequence [a] | Mw (Da) | *E. coli* 25922 MIC (µM) b |
|---|---|---|---|---|
| cTI | synthetic | KWCFRVCYRGICYRRCRG | 2303.77 | 8 |
| rcTI | VLP | **G**KWCFRVCYRGICYRRCRG**N** | 2474.93 | 8 [c] |
| LL37 | synthetic | LLGDFFRKSKEKIGKEFKRIVQRIKDFLRNLVPRTES | 4493.30 | 32 |
| rLL37 | VLP | **G**LLGDFFRKSKEKIGKEFKRIVQRIKDFLRNLVPRTES**GSHHHHHH** | 5517.33 | 32 |

[a] cTI and rcTI are joined at N- and C-termini. Additional residues present in recombinant BAPs are shown in blue

[b] *Escherichia coli* strain 25922 was grown in LB media until mid-log growth phase, then diluted to achieve a final 8 x $10^5$ CFU/mL (A600 = 0.001). Peptide treatments were prepared in LB and serially diluted into 96-well plates, which were incubated for 18 h at 37 ºC following addition of diluted bacteria. Minimum inhibitory concentration (MIC) values were determined from wells where no bacterial growth was observed (verified by measuring A600) for two biological replicates.

[c] 60.3 ± 2.8 % inhibition of growth was detected for 4 µM recombinant cTI (rcTI), compared to 17.9 ± 2.9 µM for synthetic cTI

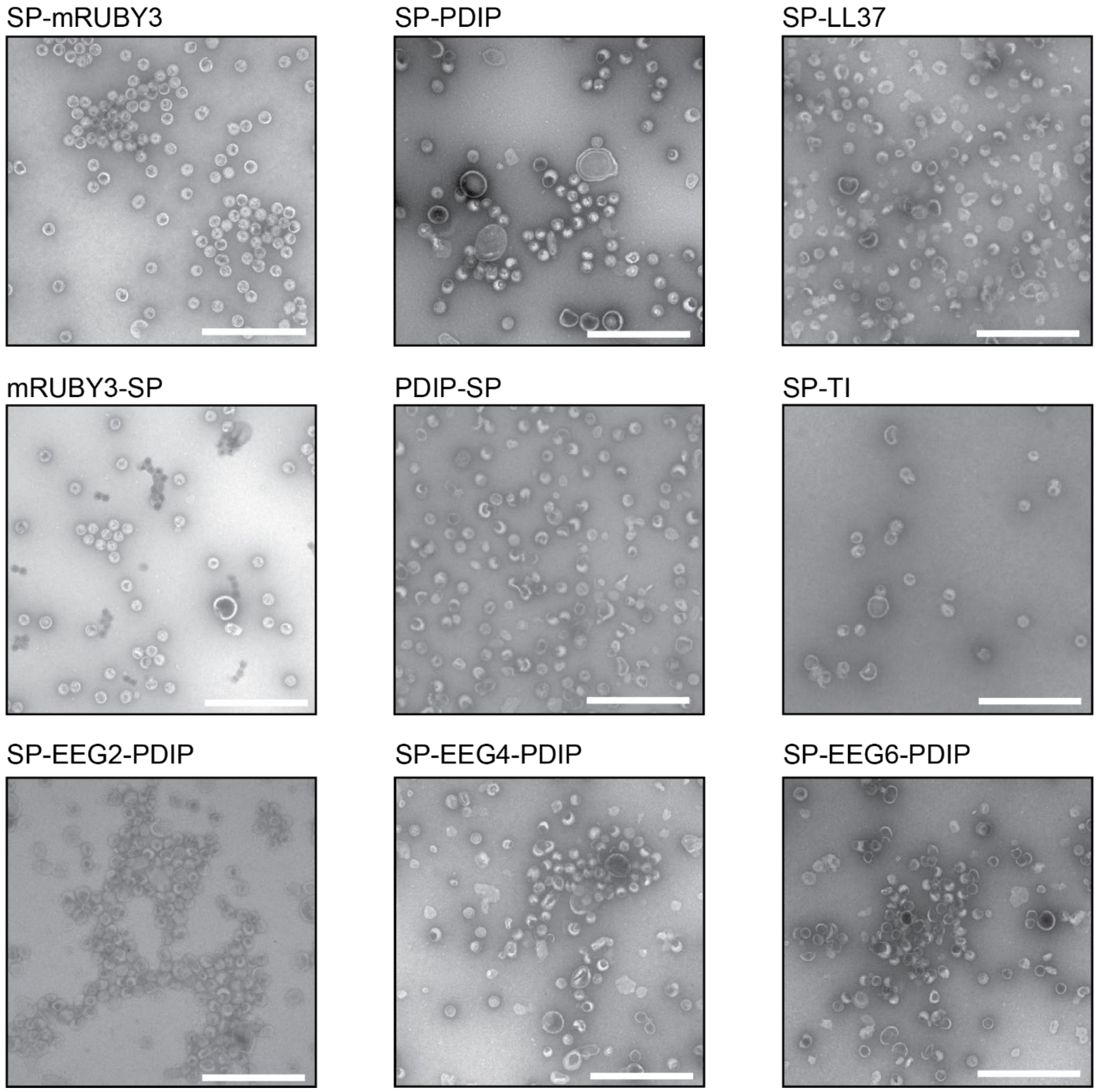


**Figure S1. Expanded TEM images of representative P22 VLPs expressed in this study.** P22 VLPs purified by ultracentrifugation were quantified using a BCA assay and 0.25 mg/mL of VLP sample was analysed by negative stain TEM on a JEOL 1010 at 80 kV. The SP fusion protein packaged within each respective P22 VLP is labelled above each image. Scale bars are 500 nm.

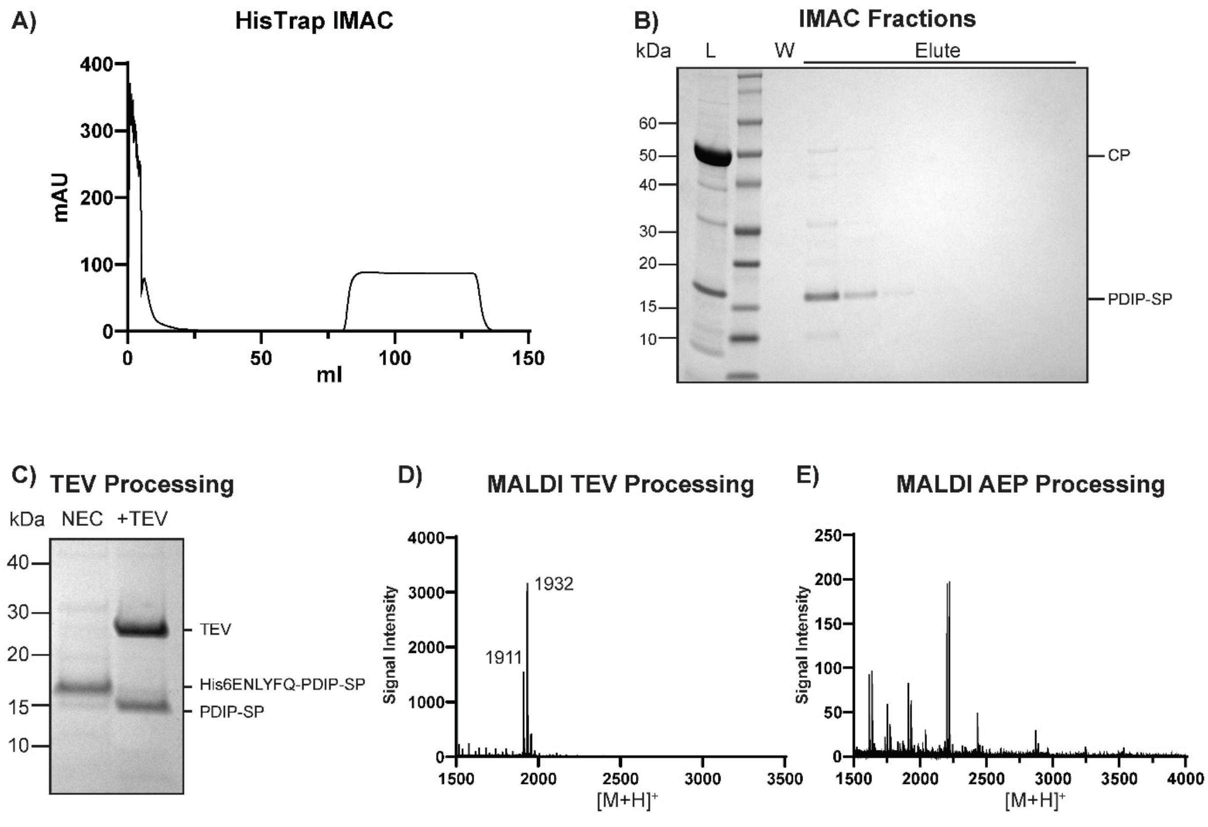


**Figure S2. Inefficient AEP processing of SP-C PDIP fusion. A)** FPLC chromatogram of IMAC purified PDIP-SP and **B)** the corresponding SDS-PAGE for the eluted PDIP-SP fractions. **C)** The SP-C orientation contains an N-terminal hexa-histidine tag, the TEV processing site and PDIP. *In vitro* processing by TEV is required to cleave the N-terminal histidine tag and liberate a free N-terminal glycine for nucleophilic attack upon AEP-mediated cyclization. The SDS-PAGE shows the molecular weight shift of the SP-C PDIP fusion upon addition of TEV (hTEV60) compared to a no enzyme control (NEC). The remaining protein is now comprised of PDIP followed by an AEP recognition sequence and the C-terminal SP (PDIP-NGL-SP). **D)** MALDI-MS spectra showing the mass of the N-terminal hexa-histidine tag and TEV recognition sequence liberated from N-terminus of PDIP-SP. **E)** A representative MALDI-MS spectra of the AEP cyclization reaction with PDIP-SP at pH 6. Various attempts at a range of time points failed to generate either cyclic or hydrolysed PDIP, which should be observed at ~3900 $[M+H]^+$.

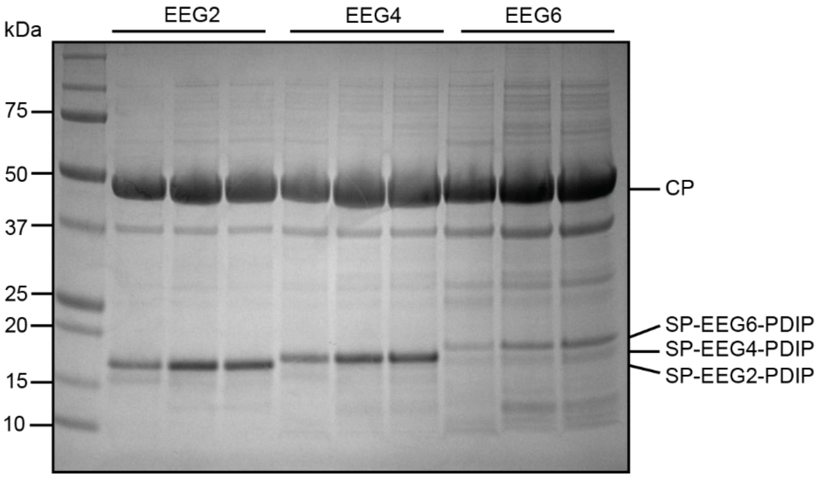


**Figure S3. SDS-PAGE gel of VLP-associated proteins from SP-(EEG)*n*-PDIP expression.** The charge effect of the anionic linker was compared by quantifying VLPs collected by ultracentrifugation using a BCA assay to normalize loading of 10 µg, then performing densitometric analysis of bands on SDS-PAGE to calculate cargo to CP ratio. Gel contains n=3 biological expression replicates for each construct.

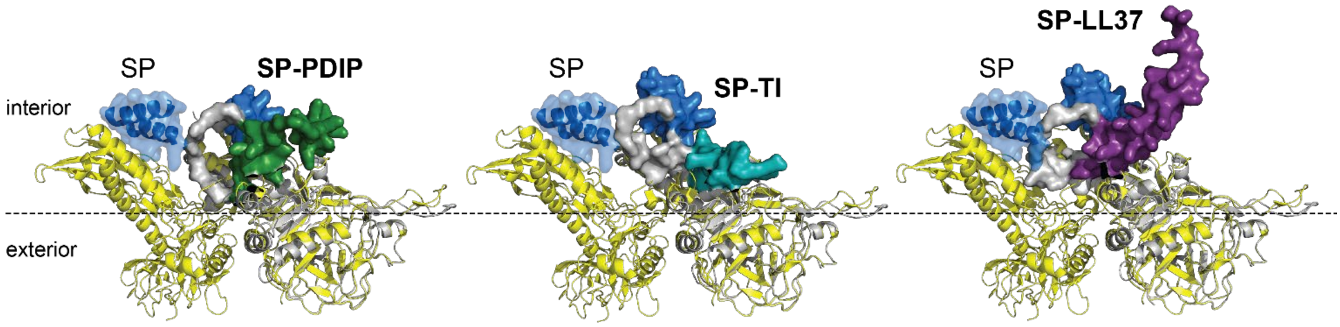


**Figure S4. Predicted interactions between SP-BAP fusions and P22 CP.** Transverse view showing CP subunits (yellow ribbon) and bound SP (blue transparent surface with ribbon) (from PDB:8I1V), overlaid with the predicted structures and interaction of CP (grey cartoon) with SP-BAPs (surface representation with SP shown in blue and BAP components shown in green for PDIP, teal for TI, and purple for LL37) (AlphaFold2-multimer with MMseqs2 on ColabFold v 1.5.5).

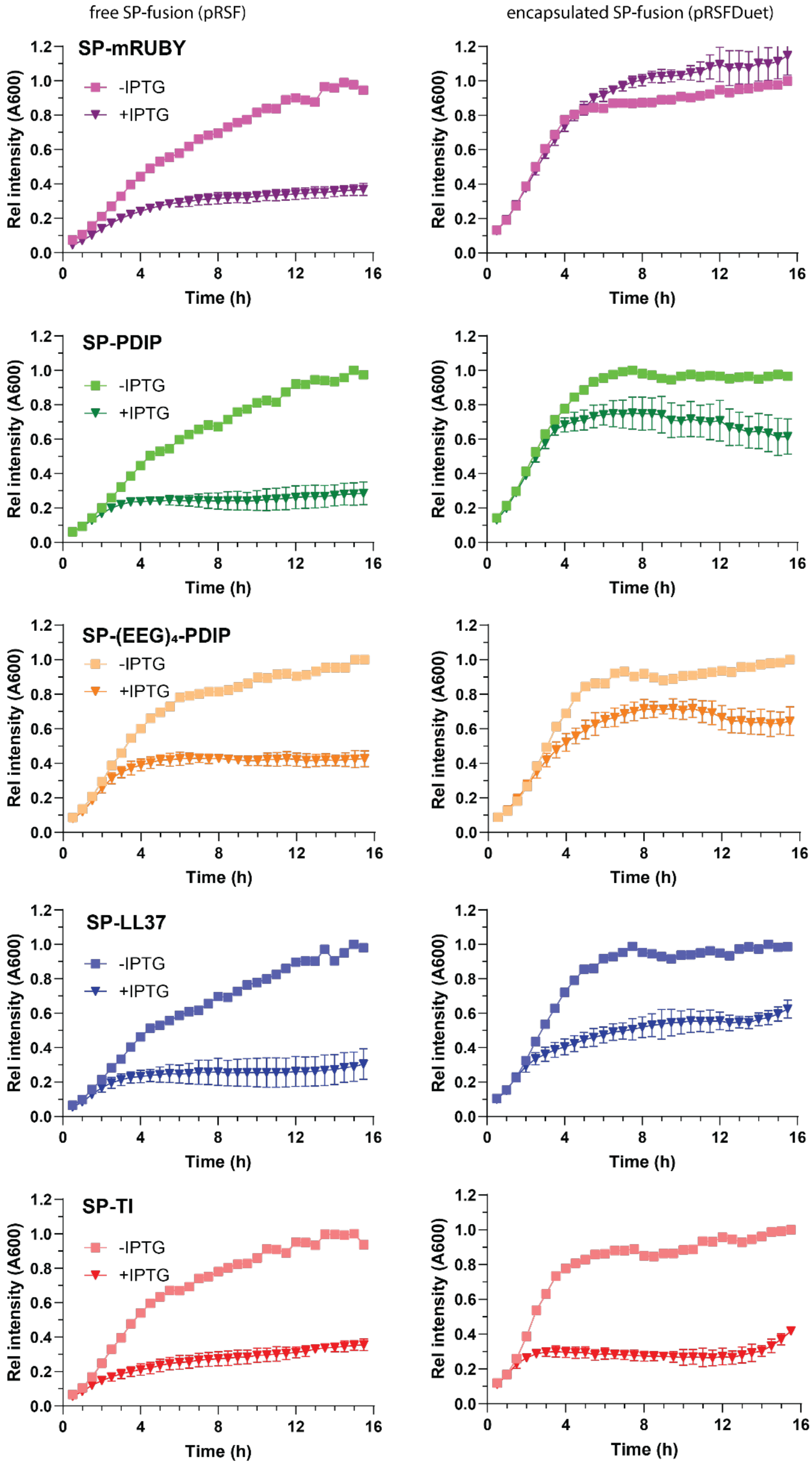


**Figure S5. Growth curves of *E. coli* expressing encapsulated (pRSFDuet) compared to free (pRSF) SP-fusions.** Growth phase cultures were prepared for three biological replicates, then diluted to OD600 of 0.1 into wells of a 96-well plate and induced with IPTG to a final concentration of 1 mM. Uninduced control wells were included to account for growth differences due to plasmid replication. The plates were incubated at 28C for 16 h with A600 recorded every 30 min. Data shown has been normalised for each construct relative to the maximum A600 achieved for the uninduced control culture. Data from induced cultures represents n = 3 biological replicates.

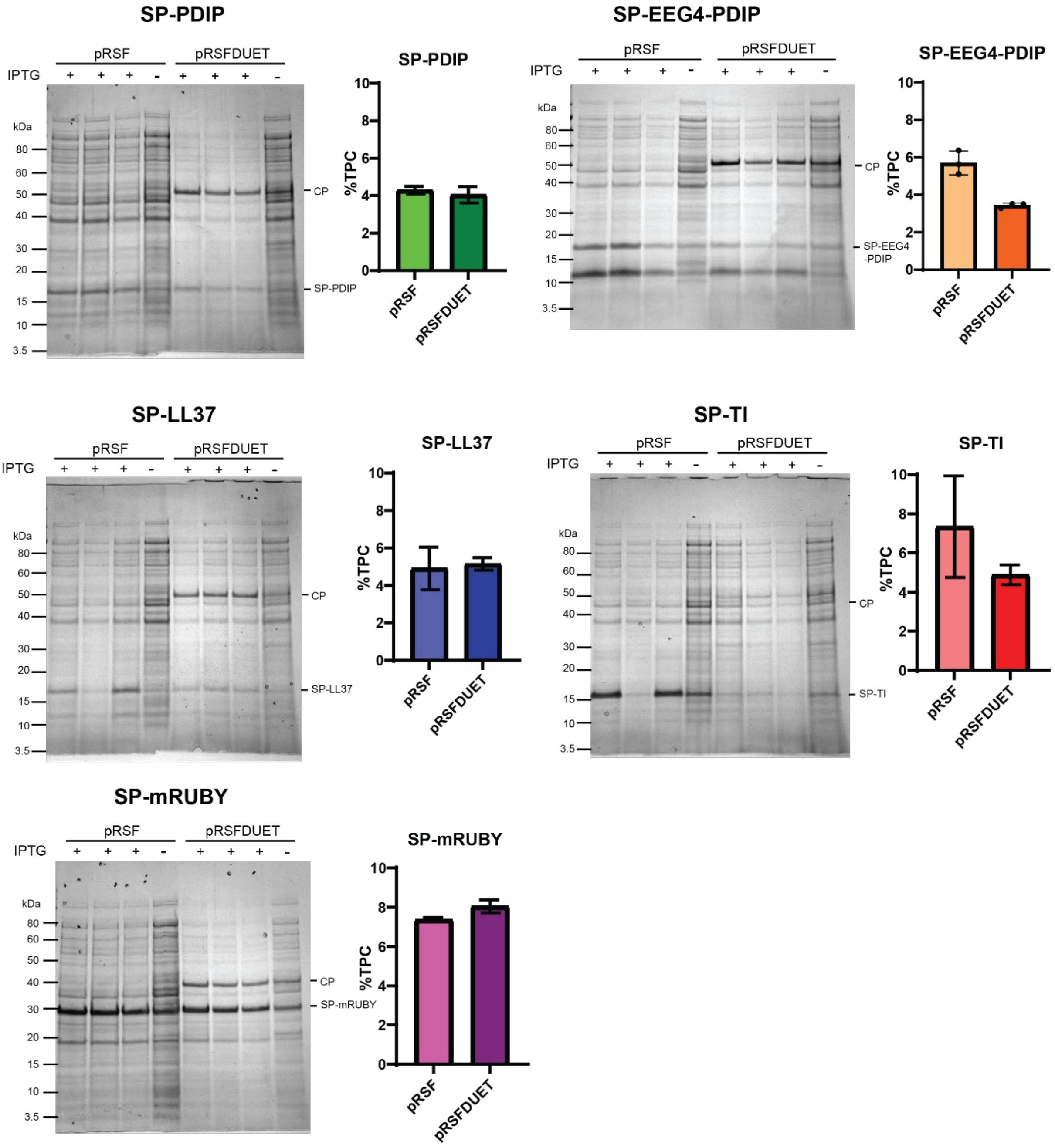


**Figure S6. Comparison of total proteins from *E. coli* expressing encapsulated (pRSFDuet) compared to free (pRSF) SP-fusions.** SDS-PAGE and the corresponding calculated yield for each SP-BAP as a percentage of total protein content (%TPC). *E. coli* total protein samples were collected following 16 h induction at 28 °C in a 96-well plate, n=1 for uninduced and n=3 for induced samples.

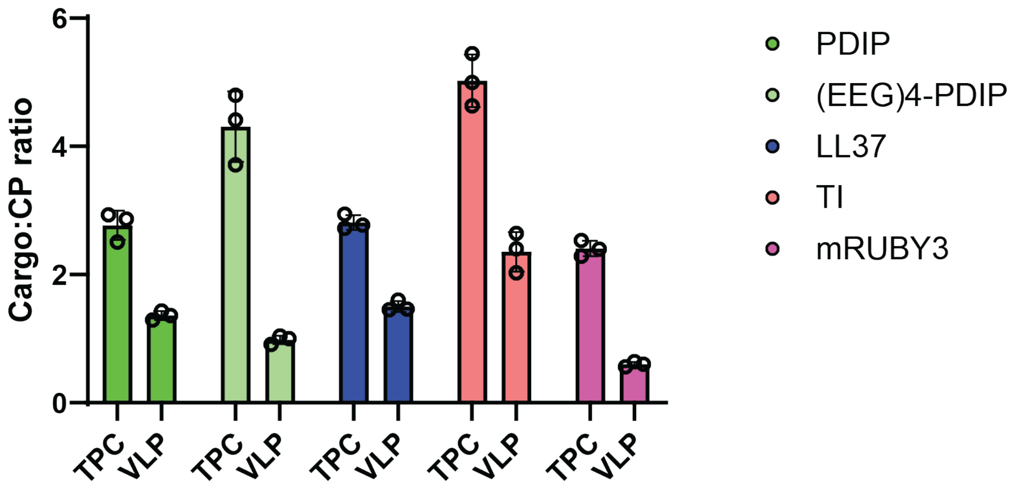


**Figure S7. Comparison of cargo to coat protein (CP) ratio for SP-fusions**. Ratios were determined from densitometric analysis of bands from SDS-PAGE gels from either total protein content of *E. coli* after 16 h induction (TPC, see Figure S6); or from virus like particles (VLPs) collected by ultracentrifugation (see Figure 2C for mRUBY and PDIP, Figure S3 for (EEG)4-PDIP, Figure 5A,B for LL37 and TI).

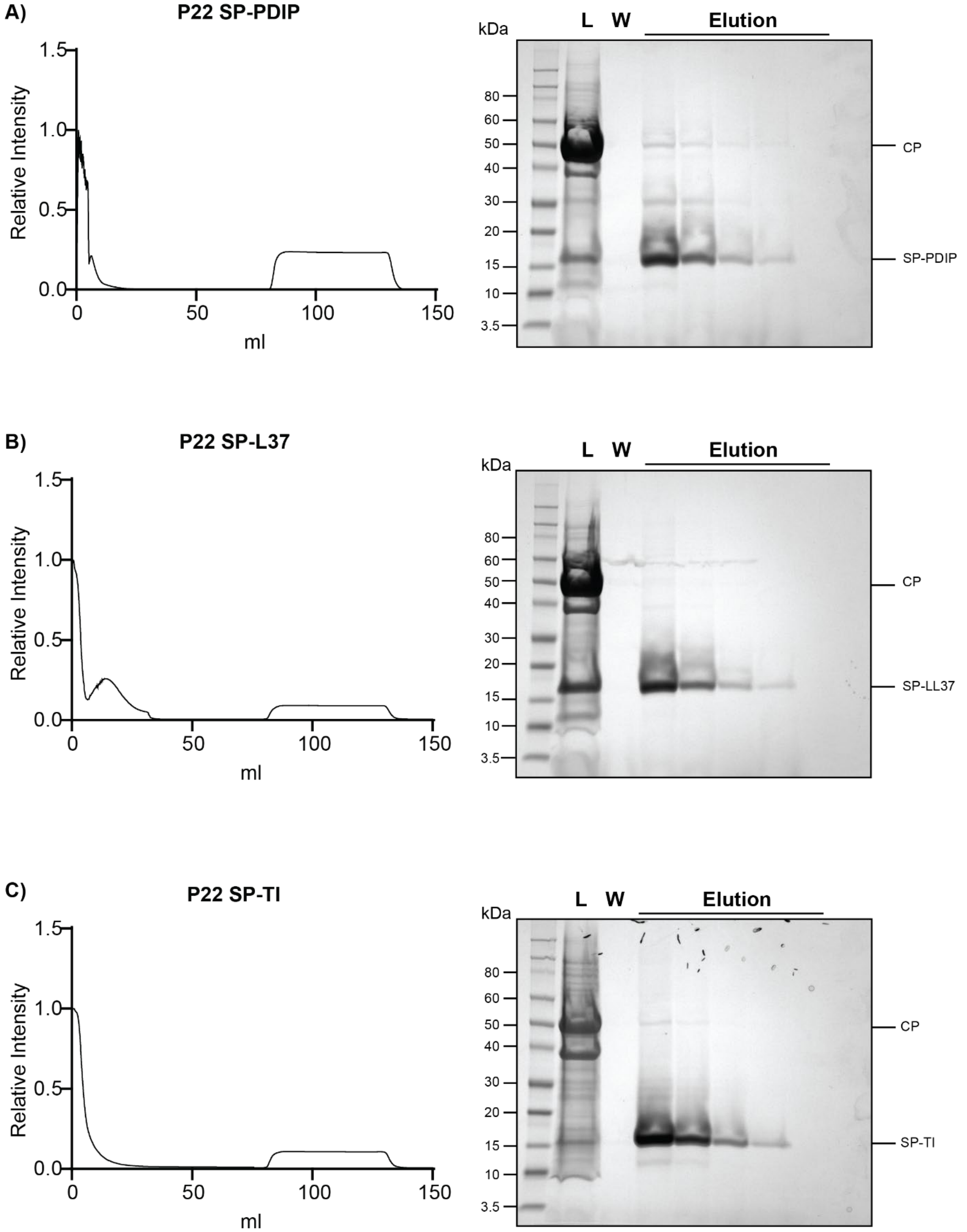


**Figure S8. Immobilized metal affinity chromatography (IMAC) and SDS-PAGE of extracted SP-BAP fusion proteins.** Purified VLPs were diluted into the chaotropic cargo extraction buffer and incubated overnight before IMAC purification using a Cytiva HisTrap column on an ÄKTA FPLC. FPLC traces and corresponding SDS-PAGE gels from IMAC purification of SP-BAPs under chaotropic conditions for **(A)** SP-PDIP, **(B)** SP-LL37, and **(C)** SP-TI. The Y-axis of FPLC chromatograms depicts a relative intensity of the mAU at 280nm absorbance, normalised to the highest absorbance value for each respective trace. Labels above SDS-PAGE gels represent an aliquot of the loaded (L) VLP sample, an aliquot from the column wash (W) and fractions from the IMAC elution.

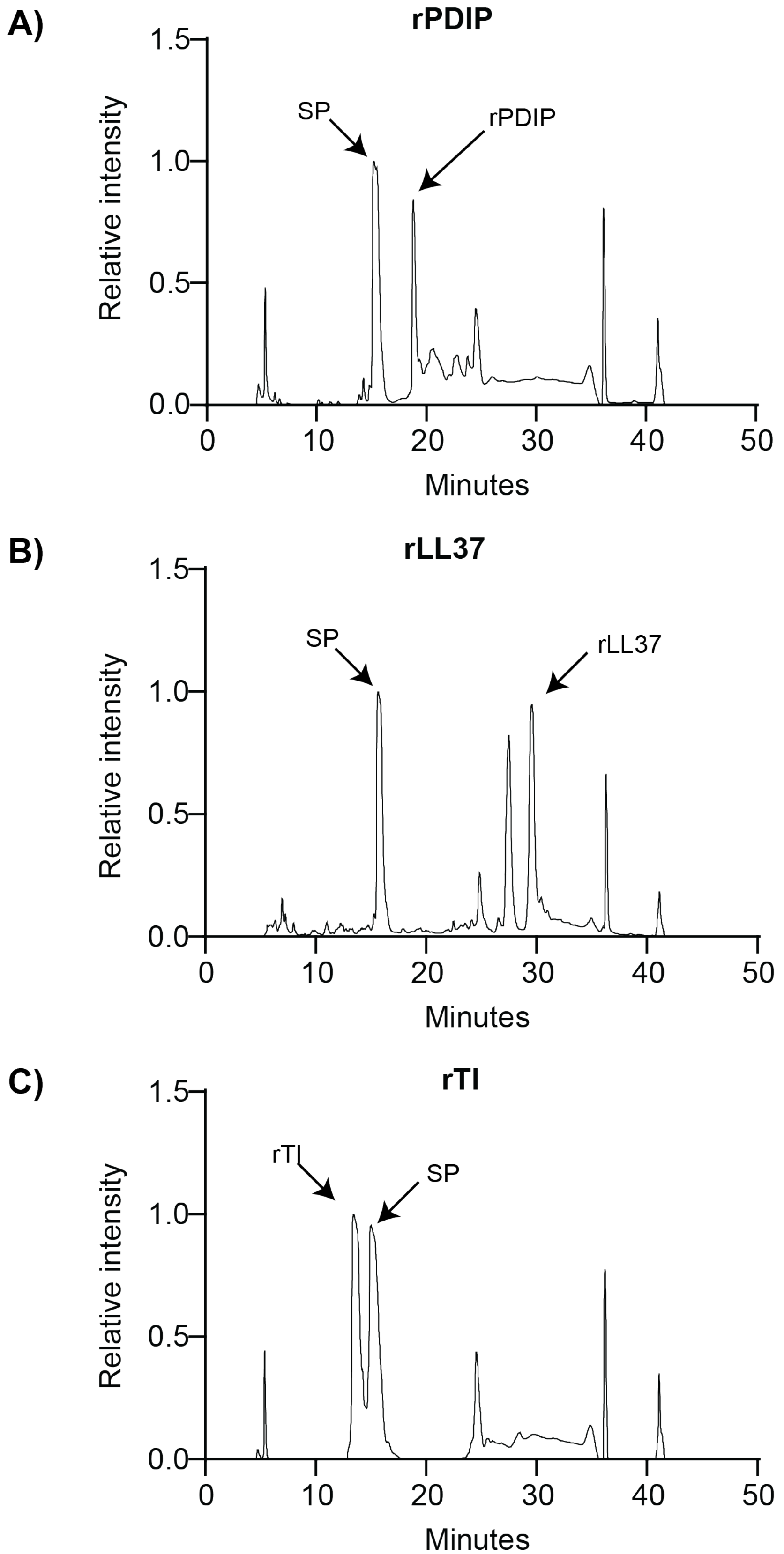


**Figure S9. HPLC traces for purification of rBAPs after TEV cleavage.** HPLC chromatogram showing relative intensity of mAU at 215nm. Arrows indicate the peaks containing respective BAPs **(A)** rPDIP, **(B)** rLL37, **(C)** rTI, cleaved from SP. Peaks at 17 and 24 min correspond to SP and TEV. The additional peak at 29 min for rLL37 is most likely a truncated form of LL37. HPLC was performed using a Phenomenex Jupiter C18 column on a Shimadzu system using a 1% gradient from solvent A (0.05% (v/v) TFA in water) to solvent B (0.05% (v/v) TFA in 90% acetonitrile).

**rPDIP**

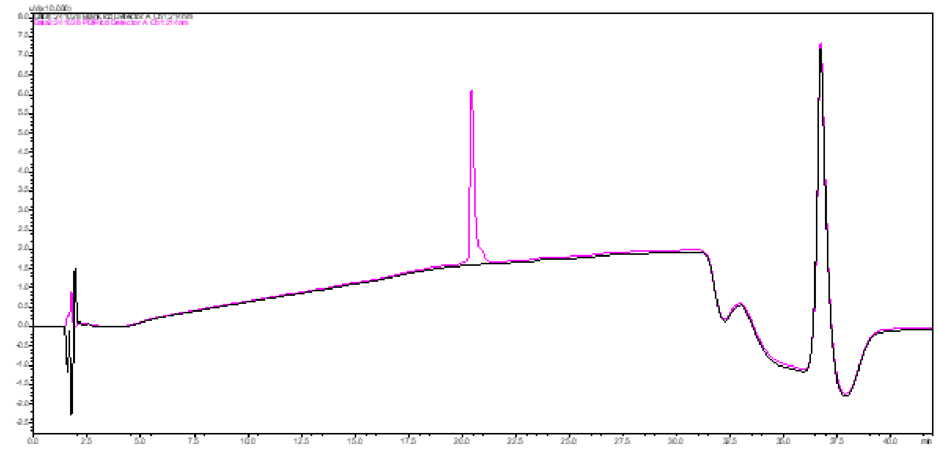

**rTI**

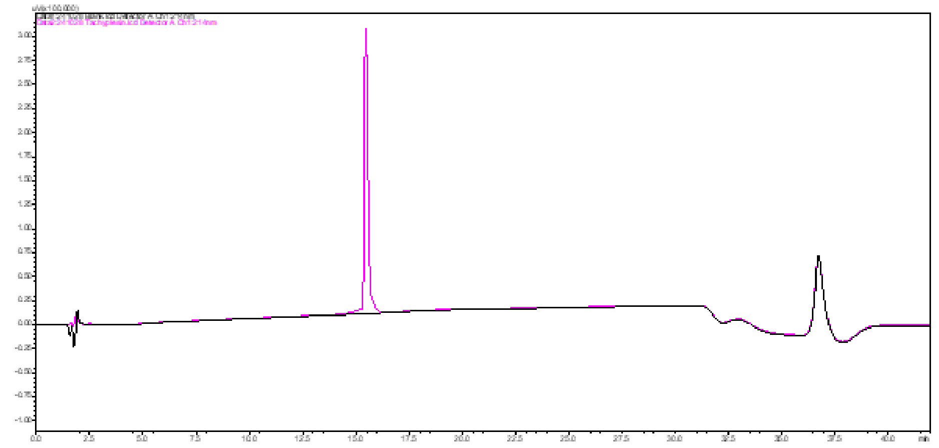

**Figure S10. UPLC trace of TEV cleaved rBAPs.** UPLC traces of purified BAPs from TEV cleavage reaction (shown in pink with solvent background in black).

**rLL37**

Calculated mass: 5517.325 Da Observed mass: 5517 Da

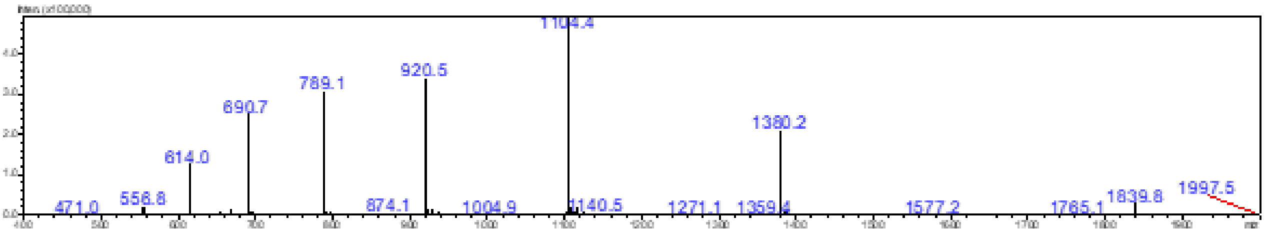


**rPDIP**

Calculated mass (oxidised): 5041.914 Da Observed mass: 5042 Da

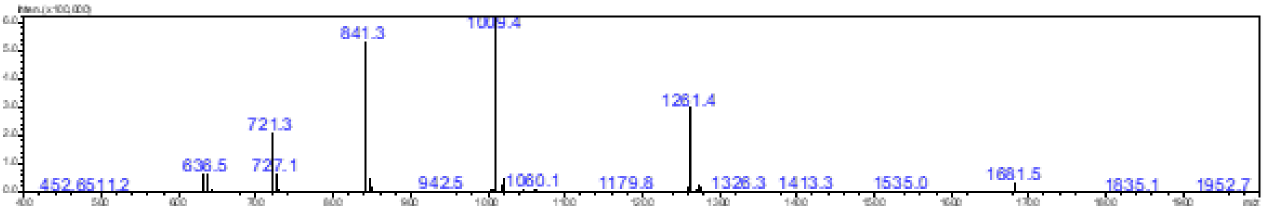


**rTI**

Calculated mass (oxidised): 3630.154 Da Observed mass: 3630 Da

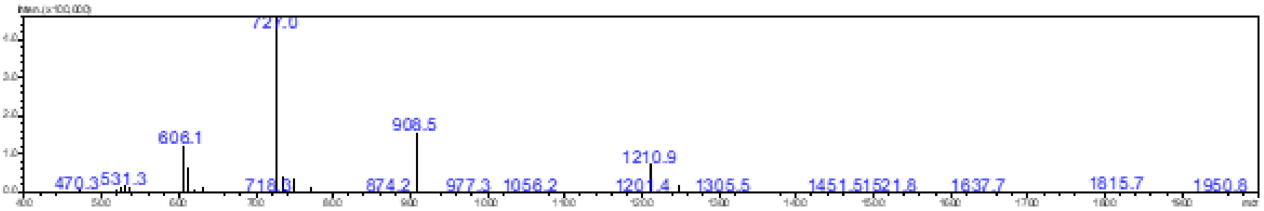


**Figure S11. ESI-MS of TEV-cleaved and HPLC-purified rBAPs.** To confirm correct cleavage by TEV, HPLC purified BAPs were assessed by ESI-MS. Calculated molecular weights for oxidised BAPs cleaved from SP are shown along with the observed masses extrapolated from the $[(m+z)/z]^{z+}$ peaks shown.

### rcTI

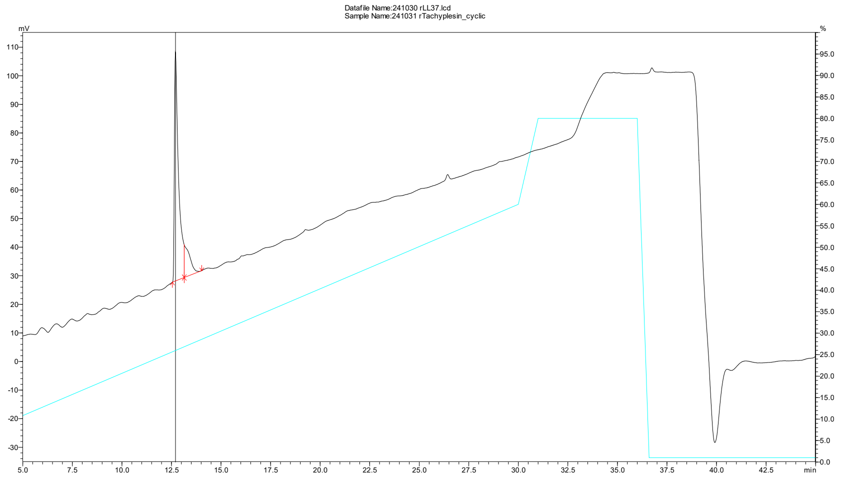


### ESI-MS main peak

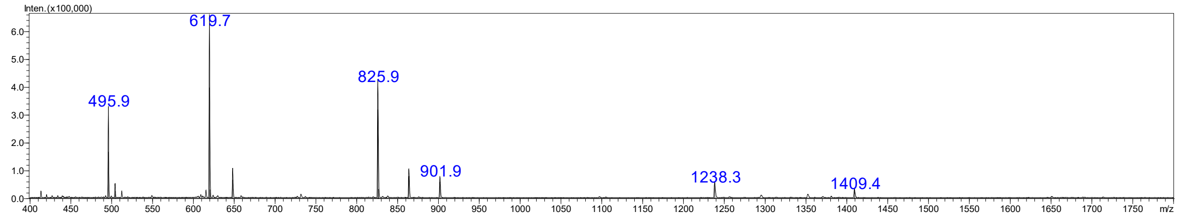


### ESI-MS trailing shoulder

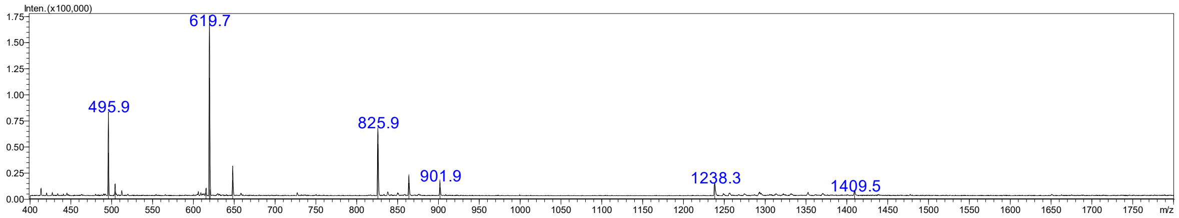


**Figure S12. LC-MS of rcTI.** LC trace of HPLC purified rcTI and corresponding ESI-MS signals $[(m+z)/z]^{z+}$ corresponding correctly to the molecular weight of cyclic, oxidised rcTI: 2474.93 Da.
